# Catalytic activity of the Bin3/MEPCE methyltransferase domain is dispensable for 7SK snRNP function in *Drosophila melanogaster*

**DOI:** 10.1101/2023.06.01.543302

**Authors:** Ryan J Palumbo, Steven D Hanes

**Affiliations:** Department of Biochemistry & Molecular Biology, SUNY Upstate Medical University 750 East Adams Street, 4283 Weiskotten Hall, Syracuse, New York, 13210

**Author notes:** **Unexpected *in vivo* results with Bin3**.

**Keywords:** Bin3, Bmc1, Bcdin3, MEPCE, Mepcea, 7SK, P-TEFb, RNA Polymerase II, Promoter-Proximal Pausing, Transcription Elongation

## Abstract

Methylphosphate Capping Enzyme (MEPCE) monomethylates the gamma phosphate at the 5’ end of the 7SK noncoding RNA, a modification thought to protect 7SK from degradation. 7SK serves as a scaffold for assembly of a snRNP complex that inhibits transcription by sequestering the positive elongation factor P-TEFb. While much is known about the biochemical activity of MEPCE *in vitro*, little is known about its functions *in vivo*, or what roles— if any—there are for regions outside the conserved methyltransferase domain. Here, we investigated the role of Bin3, the *Drosophila* ortholog of MEPCE, and its conserved functional domains in *Drosophila* development. We found that *bin3* mutant females had strongly reduced rates of egg-laying, which was rescued by genetic reduction of P-TEFb activity, suggesting that Bin3 promotes fecundity by repressing P-TEFb. *bin3* mutants also exhibited neuromuscular defects, analogous to a patient with *MEPCE* haploinsufficiency. These defects were also rescued by genetic reduction of P-TEFb activity, suggesting that Bin3 and MEPCE have conserved roles in promoting neuromuscular function by repressing P-TEFb. Unexpectedly, we found that a Bin3 catalytic mutant (Bin3 ^Y795A^) could still bind and stabilize 7SK and rescue all *bin3* mutant phenotypes, indicating that Bin3 catalytic activity is dispensable for 7SK stability and snRNP function *in vivo*. Finally, we identified a metazoan-specific motif (MSM) outside of the methyltransferase domain and generated mutant flies lacking this motif (Bin3 ^ΔMSM^). Bin3^ΔMSM^ mutant flies exhibited some—but not all—*bin3* mutant phenotypes, suggesting that the MSM is required for a 7SK-independent, tissue-specific function of Bin3.

## INTRODUCTION

Regulation of eukaryotic RNA polymerase II (RNAPII) occurs at multiple stages of the transcription cycle, including RNAPII recruitment to promoters by enhancer binding proteins, the establishment of the pre-initiation complex, and promoter release. In higher eukaryotes an additional mechanism, promoter-proximal pausing, has been shown to be critical for RNAPII regulation in development and in response to signaling events (Margaritis and Holstege 2008; Gilmour 2009; Chiba *et al*. 2010; Levine 2011; Li and Gilmour 2011; Nechaev and Adelman 2011; Adelman and Lis 2012; Zhou *et al*. 2012; Gaertner and Zeitlinger 2014; Jonkers and Lis 2015; Liu *et al*. 2015; Mayer*et al*. 2017; Chen *et al*. 2018; Core and Adelman 2019; Dollinger and Gilmour 2021; Gonzalez *et al*. 2021; Abuhashem *et al*. 2022). Pausing occurs when RNAPII stalls after transcribing approximately 30-60 nucleotides, and is induced by the binding of negative elongation factors (Yamaguchi *et al*. 1999; Wang *et al*. 2010) and by hypo-phosphorylation of serine 2 (Ser2) in the heptad repeat (YSPTSPS) of the carboxy-terminal domain of the large subunit of RNAPII (reviewed in Brookes and Pombo 2009). Promoter proximal pausing is relieved by the kinase activity of Positive Transcription Elongation Factor b (P-TEFb; Marshall and Price 1995), which is a heterodimer composed of Cyclin-dependent kinase 9 (Cdk9; Marshall *et al*. 1996) and the regulatory cyclin, Cyclin T (CycT; Peng *et al*. 1998). P-TEFb phosphorylates negative elongation factors (Wada *et al*. 1998; Cheng and Price 2007) and Ser2 (Zhou *et al*. 2000; Ramanathan *et al*. 2001; Shim *et al*. 2002; Ni *et al*. 2004), which allows RNAPII to enter productive elongation (Ni *et al*. 2008).

To maintain a balance between transcription elongation and pausing, P-TEFb activity is highly regulated; without this regulation, P-TEFb becomes “hyper-activated” and induces aberrant transcription elongation (Schneeberger *et al*. 2019). Key to this regulation is the 7SK snRNP, which sequesters (and thereby represses) P-TEFb (Nguyen *et al*. 2001; Yang *et al*. 2001). The “core” 7SK snRNP is composed of the 7SK noncoding RNA, which acts as a scaffold for the proteins Methylphosphate Capping Enzyme (MEPCE) and La-Related Protein 7 (LARP7) (Krueger *et al*. 2008; Barboric *et al*. 2009; Xue *et al*. 2010; Muniz *et al*. 2013; Brogie and Price 2017; Yang *et al*. 2022). MEPCE and LARP7 protect the 5’ and 3’ ends of 7SK from exoribonucleolytic degradation, respectively (Jeronimo *et al*. 2007; Markert *et al*. 2008; Barboric *et al*. 2009; Xue *et al*. 2010; Muniz *et al*. 2013). After formation of the core snRNP, 7SK RNA is reversibly bound by a homodimer of Hexamethylene Bis-Acetamide Inducible Protein (HEXIM) (Blazek *et al*. 2005; Li *et al*. 2005). Binding of HEXIM to 7SK induces a conformational change that allows HEXIM to bind to P-TEFb (Michels *et al*. 2004), functionally sequestering and inactivating P-TEFb (Yik *et al*. 2004). Upon cellular stress and other cellular signals (reviewed in Liu *et al*. 2015), binding of accessory proteins to 7SK (Herreweghe *et al*. 2007; Barrandon *et al*. 2007; Krueger *et al*. 2008; Lemieux *et al*. 2015; Bugai *et al*. 2019) and a concomitant conformational change in the secondary structure of 7SK RNA (Krueger *et al*. 2010; Yang *et al*. 2022) releases P-TEFb from the repressive 7SK snRNP. P-TEFb is then able to relieve pausing to enable transcription elongation.

MEPCE is the long sought-after RNA methyltransferase that uses the methyl donor S-adenosyl-methionine (SAM) to add a monomethyl cap to the gamma phosphate on the 5’ end of 7SK (Gupta *et al*. 1990a; Shumyatsky *et al*. 1990; Shimba and Reddy 1994), an RNA polymerase III transcript (Zieve and Penman 1976; Murphy *et al*. 1986, 1987, 1989; Shumyatsky *et al*. 1990). MEPCE is human ortholog of Bin3, a *Drosophila* protein that is the founding member of the Bin3/MEPCE/Bmc1 family of RNA methyltransferases (Zhu and Hanes 2000; Jeronimo *et al*. 2007; Barboric *et al*. 2009; Cosgrove *et al*. 2012; Páez-Moscoso *et al*. 2022; Porat *et al*. 2022). We originally discovered Bin3 as an interacting partner of the Bicoid homeodomain protein (Zhu and Hanes 2000). Bicoid acts as both an activator of zygotic gene transcription, and a translational repressor of maternal *caudal* mRNA during embryogenesis; both functions are critical for anterior-posterior patterning (Dubnau and Struhl 1996; Rivera-Pomar *et al*. 1996; Chan and Struhl 1997; Niessing *et al*. 1999, 2000, 2002; Cho *et al*. 2005). We had hypothesized that Bin3 was an arginine methyltransferase that methylated Bicoid to alter its function, transitioning Bicoid from being a transcriptional activator to a translational repressor, or vice versa. However, identification and characterization of MEPCE in human cells (originally called BCDIN3) demonstrated that it is actually an RNA methyltransferase that targets 7SK to form a snRNP that represses transcription (Jeronimo *et al*. 2007). We later showed that early in*Drosophila* embryogenesis, Bin3 stabilizes 7SK to form a novel snRNP containing Bin3 and Bicoid (and potentially other translational factors) that represses translation of maternal *caudal* mRNA at the anterior of the embryo (Singh *et al*. 2011). It is important to note that Bicoid is an insect-specific protein, and therefore, the translational repression function of Bin3 is also likely to be insect-specific. The canonical role of Bin3/MEPCE in regulating transcription elongation, and the importance of this regulation to normal development, has not been extensively studied in *Drosophila* or in other model organisms (see below).

Much of what we know about the 7SK RNA binding and capping functions of MEPCE comes from *in vitro* or cell culture-based studies focusing on the methyltransferase activity of MEPCE and the stepwise assembly of the 7SK snRNP. The monomethyl cap is believed to protect the 5’ end of 7SK from exoribonucleolytic degradation (Shumyatsky *et al*. 1993). However, MEPCE is peculiar in that unlike most enzymes, it remains bound to the product of the reaction it catalyzes, calling into question the essentiality of methyl capping. MEPCE that is constitutively bound to capped 7SK facilitates the stable binding of LARP7 (Xue *et al*. 2010) to form the core 7SK snRNP (Muniz *et al*. 2013). Several residues in the active site have been identified as being essential for binding to SAM and/or 7SK (Xue *et al*. 2010; Shelton *et al*. 2018), and for methyltransferase activity (Yang *et al*. 2019). Pre-association of MEPCE with 7SK was found to enhance the recruitment of LARP7 to 7SK and to MEPCE itself (Xue *et al*. 2010). However, whether methyltransferase activity is required *in vivo* for 7SK binding and stability, and 7SK snRNP function, has never explicitly been tested. Moreover, these studies have focused primarily on the function of the methyltransferase domain. During evolution, MEPCE orthologs have acquired extensive protein sequence flanking the methyltransferase domain, the conservation and function(s) of which have not been explored.

Underscoring the need to understand the function of MEPCE *in vivo*, there is a neurodevelopmental disorder that is caused by heterozygosity for a nonsense mutation in *MEPCE* that results in P-TEFb hyper-activation (Schneeberger *et al*. 2019). However, our ability to study the *in vivo* function of MEPCE and its orthologs is limited by the fact that MEPCE function appears to be essential in vertebrate model organisms. For example, transfecting HeLa cells with siRNAs to destabilize 7SK (which phenocopies the destabilization of 7SK in the absence of MEPCE function) results in apoptosis (Haaland *et al*. 2005); treating zebrafish embryos with morpholino oligonucleotides targeting *mepcea* (the zebrafish ortholog of *MEPCE* ; originally named *bcdin3*) is embryonic lethal (Barboric *et al*. 2009), precluding the ability to study Mepcea function in adult zebrafish; and deleting 7SK in mice (which also phenocopies the destabilization of 7SK in the absence of MEPCE function) is lethal (Xu *et al*. 2022), precluding the ability to study MEPCE function in adult mice. However, deleting *bin3* (the *Drosophila* ortholog of *MEPCE*) does not affect the viability of adult *Drosophila* (Singh *et al*. 2011). Therefore, *Drosophila* Bin3 provides a model to study the *in vivo* functions of MEPCE in metazoan development.

Here, we investigated the role of Bin3 in *Drosophila* development. We found that Bin3 is required for fecundity and has a conserved role in promoting neuromuscular function, by repressing P-TEFb. Unexpectedly, we found that Bin3 catalytic activity is dispensable for 7SK stability and snRNP function *in vivo*, challenging the paradigm that methyl capping is essential for 7SK stability. Finally, we identified a metazoan-specific motif (MSM) outside of the methyltransferase domain that confers a 7SK-independent, tissue-specific function to Bin3. These studies illustrate the importance of establishing a model system in which to study this important and conserved enzyme.

## MATERIALS AND METHODS

### Fly stocks and husbandry

Fly stocks from the Bloomington *Drosophila* Stock Center (BDSC) used in this study are listed in **Table S1**. Fly stocks we constructed for this study are listed in **Table S2**. The genotypes of maternal and paternal flies, and the genotypes of their progeny used for experiments, and the figures in which those experiments appear, are listed in **Table S3**. Fly were reared at 25°C and 60-80% humidity on a 12-hour light/dark cycle, on either Nutri-Fly MF food (Genesee Scientific) with ∼0.064 M propionic acid (Sigma Aldrich, P5561; or Apex Bioresearch Products, 20-271), or food made based on the Nutri-Fly MF formula: 25 g/L Inactive Dry Yeast (Genesee Scientific), 89.5 g/L Dry Molasses (Genesee Scientific), 57 g/L Fly Stuff Yellow Cornmeal (Genesee Scientific), 5.84 g/L Nutri-Fly *Drosophila* Agar, Gelidium (Genesee Scientific), ∼0.064 M propionic acid (Apex Bioresearch Products).

### *bin3* mutants

We previously described two excision alleles that remove some (*bin3^4-7^*) or all (*bin3^2-7^*) of the *bin3* coding exons (Parks *et al*. 2004; Singh *et al*. 2011). Both alleles were made in the same background, and when placed in *trans* to produce *bin3^4-7^/bin3^2-7^* mutants, incurred background effects likely due to homozygosity for recessive second-site mutations. To produce *bin3* mutants that avoid these confounding effects, we crossed flies heterozygous for *bin3^2-7^* (**Fig. 1a**) with flies heterozygous for a deficiency uncovering *bin3* (*Df*, **Fig. 1a**), which is in a different genetic background than *bin3^2-7^* (Ryder *et al*. 2004).

**Figure 1.**
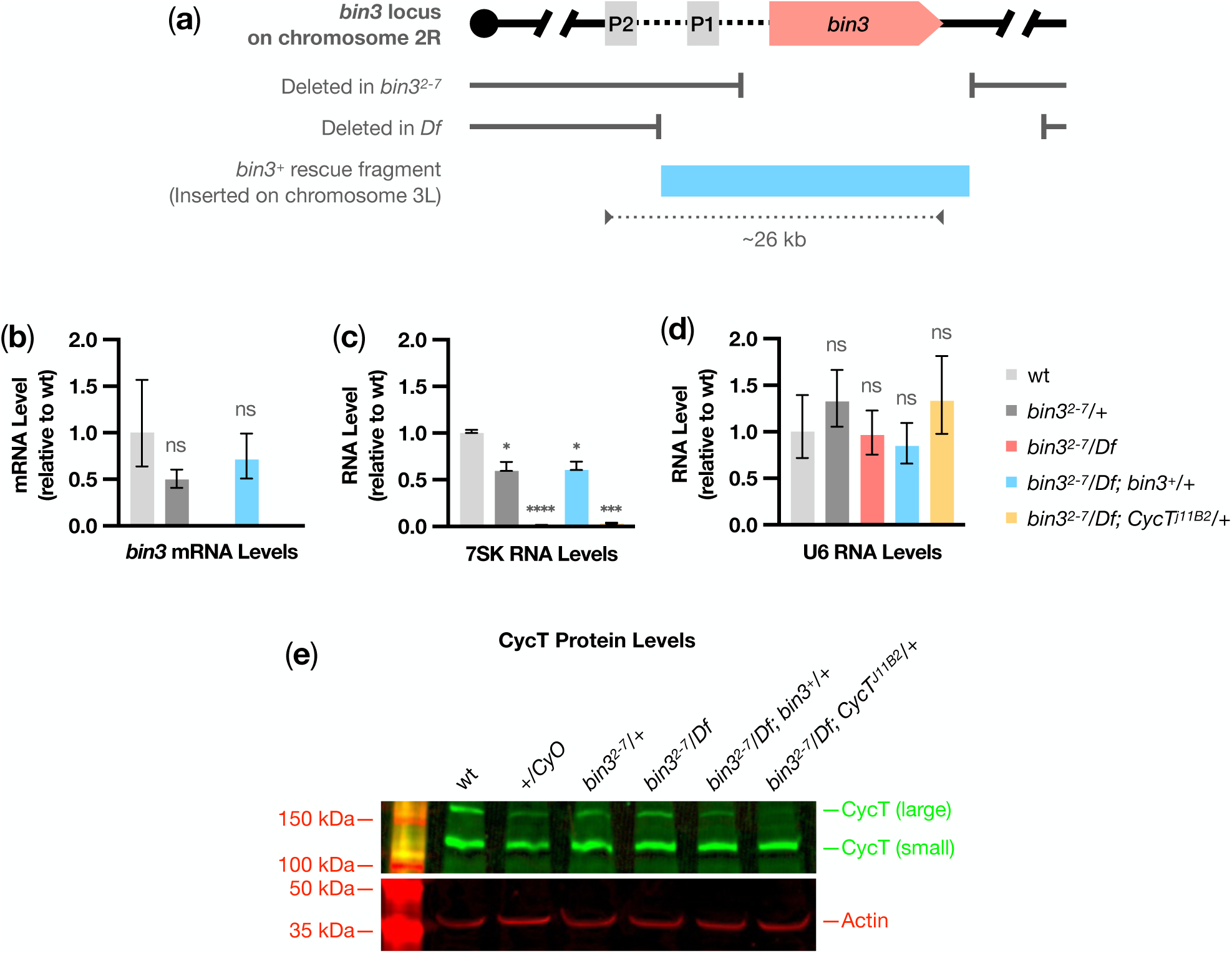
Molecular characterization of *bin3* mutant and rescue flies. **(a)** Graphic depicting the ∼26 kb *bin3* locus on chromosome 2R (cytogenetic bands 42A13-42A14). Two promoters, P1 and P2 (grey), drive expression of *bin3* (Zhu and Hanes 2000; Singh *et al*. 2011). The *bin3* ORF (salmon) comprises 6 coding exons. *bin3^Δ^* refers to the *bin3^2-7^* excision allele created by FLP/FRT-mediated recombination between transposon insertions *PBac{RB}bin3^e02231^* and *P{XP}d02161* (Parks *et al*. 2004; Singh *et al*. 2011). *Df* refers to deficiency *Df(2R)ED1612*, which deletes ∼829 kb of chromosome 2R, including promoter P1 and all *bin3* coding sequence (Ryder *et al*. 2004). *bin3^Δ^*and *Df* in *trans* produce *bin3* mutant flies. *bin3^+^* (sky blue) is a transgenic insertion of a BAC comprising ∼20 kb of chromosome 2R, including promoter P1 and all *bin3* coding sequence. In *bin3^2-7^/Df* flies, this insertion rescues *bin3* mutant phenotypes. **(b-d)** Bar graphs showing the average amount of *bin3* mRNA (b), 7SK snRNA (c), and U6 snRNA (d) in total RNA extracted from ovaries of females of the indicated genotypes, for three biological replicates. Note that *bin3* mRNA was below the level of detection in *bin3^2-7^/Df* and *bin3^2-7^/Df; CycT^j11B2^/+* ovaries for 2 of the three biological replicates. RNA levels were normalized to *rp49* mRNA and made relative to wt using CFX Maestro 2.0 (version 5.2.008.0222, Bio-Rad). Error bars represent the standard error of the mean. Statistical significance was calculated by unpaired t-test using CFX Maestro 2.0 (version 5.2.008.0222, Bio-Rad). *P* values of <.05 (*), <0.001 (***), <.0001 (****), and not significant (ns) are indicated. **(e)** Western blot for CycT (Nguyen *et al*. 2012) and Actin in extracts prepared from ovaries of females of the indicated genotypes. The *P{lacW}j11B2^jllB2^*insertion (Spradling *et al*. 1999) into one of the two different *CycT* 5’-UTRs reduces expression of the large isoform of CycT.

### Generation of transgenic *bin3^+^* rescue flies

We previously showed that Bin3 is required for embryonic viability by demonstrating a *bin3* cDNA under control of the maternal *Hsp83* promoter rescued the embryonic lethality of eggs laid by *bin3* mutant females (Singh *et al*. 2011). To examine the role of Bin3 more broadly, we needed a rescue approach that would faithfully express *bin3* under control of its own regulatory sequences in all *bin3*-expressing tissues, not just in the ovary. To do this, we used BAC clone CH322-154G8 (BACPAC Genomics) that comprises a 22.065 kb fragment of chromosome 2R, containing the *bin3* P1 promoter (Singh *et al*. 2011), and 3 of the 4 predicted mRNA isoforms of *bin3* (*bin3-RA*, *bin3-RC*, and *bin3-RD*). This BAC was inserted into the *attP* landing site on chromosome 3R in BDSC stock #9744 (*y^1^ w^1118^; PBac{y^+^-attP-9A}VK00027*) by BestGene, Inc.

### Generation of the pUAS-DSCP-XbaI-attB vector via NEBuilder HiFi DNA Assembly

High-copy plasmids pNP (Qiao *et al*. 2018; Wang *et al*. 2019) and pUASP-attB (Drosophila Genomics Resource Center Stock 1358; https://dgrc.bio.indiana.edu//stock/1358; RRID:DGRC_1358; Takeo *et al*. 2012) were purified from bacterial cells grown to confluence at 37°C in LB + 100 µg/ml ampicillin using the Monarch Plasmid Miniprep Kit T1010L from New England BioLabs, Inc. (NEB). To create pUAS-DSCP-XbaI-attB (the vector into which *bin3* and 3xHA sequences would be cloned, see below), the 10XUAS-DSCP promoter was amplified from plasmid pNP with Q5 High-Fidelity DNA Polymerase (NEB) using primers UP0394 and UP0395, and inserted into pUASP-attB that had been digested with *Xho*I (NEB) and *Pst*I (NEB), via NEBuilder HiFi DNA Assembly (NEB.). NEBuilder HiFi DNA Assembly destroyed the *Xho*I site, and partially eliminated one of the two GAGA enhancers in pUASp-attB.

### Ovarian RNA Purification and Reverse Transcription for NEBuilder HiFi DNA Assembly

one to three-day old *w^1118^* (BDSC #5905) females were incubated with males in yeasted vials overnight. Five pairs of ovaries were dissected from two to four-day old females in ice-cold 1X PBS, and flash-frozen in liquid nitrogen. Total RNA was extracted using the Monarch Total RNA Miniprep Kit (NEB), according to the manufacturer’s instructions for RNA purification from “Tissues”. Total RNA (1 µg) was reverse transcribed using ProtoScript II First Strand cDNA Synthesis Kit (NEB, E6560S) with primer UP0118, which anneals to the very end of *bin3* coding sequence (CDS). See **Table S4** for primer sequence.

### Purification of BAC clone CH322-154G8 for NEBuilder HiFi DNA Assembly

EPI300 bacterial cells (Lucigen) harboring the BAC clone CH322-154G8 (BACPAC Genomics) were grown to confluence at 37°C in LB + 12.5 µg/ml chloramphenicol, and plasmid replication was induced by diluting 1:10 in LB + 12.5 µl/ml chloramphenicol and 1X CopyControl Induction Solution (Lucigen). The BAC was purified purified using the Monarch Plasmid Miniprep Kit (NEB).

### Generation of *UAS-DSCP-3xHA-bin3* transgenic flies via NEBuilder HiFi DNA Assembly

To generate plasmid *pUAS-DSCP-3xHA-bin3-attB*, which expresses Bin3 C-terminally tagged with 3xHA, under UAS control: Fragment #1, comprising the *bin3-RA* 5’-UTR to the end of the *bin3* CDS, was amplified from *bin3* cDNA using Q5 High-Fidelity DNA Polymerase, and primers UP0423 and UP0414.

The 5’ end of Fragment #1 is homologous to sequences immediately upstream of the *Xba*I site in pUAS-DSCP-XbaI-attB; the 3’ end of Fragment #1 is homologous to 5’ end of a synthetic dsDNA fragment comprising Gly-Gly-Gly-Ser-3xHA (G _4_S-3xHA; Integrated DNA Technologies) codon-optimized for *Drosophila* with the IDT Codon Optimization Tool (Integrated DNA Technologies). Fragment #2, comprising the entire *bin3-RA* 3’-UTR, was amplified from the BAC clone CH322-154G8 using Q5 High-Fidelity DNA Polymerase, and primers UP0415 and UP0424. The 5’ end of Fragment #2 is homologous to the 3’ end of the G_4_S-3xHA fragment; the 3’ end of Fragment #2 is homologous to sequences immediately downstream of the *Xba*I site in pUAS-DSCP-XbaI-attB. Fragment #1, G _4_S-3xHA, and Fragment #2 were assembled in this order into the *Xba*I site of pUAS-DSCP-XbaI-attB via NEBuilder HiFi DNA Assembly. The *Xba*I site was destroyed.

To generate plasmid *pUAS-DSCP-3xHA-bin3 ^Y795A^-attB*, which expresses catalytically-dead Bin3 ^Y795A^, C-terminally tagged with 3xHA, under UAS control: Fragment #3, comprising the *bin3-RA* 5’-UTR to 11 nt downstream of the Y795 codon, was amplified from *bin3* cDNA using Q5 High-Fidelity DNA Polymerase, and primer UP0423 and the mutagenic primer UP0419, which introduces a Y795A point mutation. The 5’ end of Fragment #3 is homologous to sequences immediately upstream of the *Xba*I site in pUAS-DSCP-XbaI-attB; the 3’ end of Fragment #3 is homologous to the 5’ end of Fragment #4. Fragment #4, comprising 11 nt upstream of the Y795 codon to the end of the *bin3-RA* CDS was amplified from cDNA using Q5 High-Fidelity DNA Polymerase, and mutagenic primer UP0420, which introduces a Y795A point mutation, and primer UP0414. The 5’ end of Fragment #4 is homologous to the 3’ end of Fragment #3; the 3’ end of Fragment #4 is homologous to 5’ end of the G _4_S-3xHA fragment. Fragment #3, Fragment #4, G _4_S-3xHA, and Fragment #2 (see above) were assembled in this order into the *Xba*I site of pUAS-DSCP-XbaI-attB via NEBuilder HiFi DNA Assembly. The *Xba*I site was destroyed.

To generate plasmid *pUAS-DSCP-3xHA-bin3 ^ΔMSM^-attB*, which expresses Bin3^ΔMSM^, C-terminally tagged with 3xHA, under UAS control: Fragment #5, comprising the *bin3-RA* 5’-UTR to 12 nt downstream of the L326 codon, was amplified from *bin3* cDNA using Q5 High-Fidelity DNA Polymerase, and primer UP0423 and the mutagenic primer UP0417, which eliminates codons 311-326. The 5’ end of Fragment #5 is homologous to sequences immediately upstream of the *Xba*I site in pUAS-DSCP-XbaI-attB; the 3’ end of Fragment #5 is homologous to the 5’ end of Fragment #6. Fragment #6, comprising 12 nt upstream of the F311 codon to the end of the*bin3-RA* CDS was amplified from cDNA using Q5 High-Fidelity DNA Polymerase and mutagenic primer UP0418, which eliminates codons 311-326, and primer UP0414. The 5’ end of Fragment #6 is homologous to the 3’ end of Fragment #5; the 3’ end of Fragment #6 is homologous to 5’ end of the G _4_S-3xHA fragment. Fragment #5, Fragment #6, G _4_S-3xHA, and Fragment #2 (see above) were assembled in this order into the *Xba*I site of pUAS-DSCP-XbaI-attB via NEBuilder HiFi DNA Assembly. The *Xba*I site was destroyed.

See **Table S4** for primer sequences and the sequence of the G_4_S-3xHA synthetic dsDNA fragment used to make these 3 constructs. Sequencing of each construct revealed that all UTRs corresponded to the *bin3-RA* mRNA isoform of *bin3*. Interestingly, each construct had an in-frame insertion of an Ala residue between L573 and V574, when mapped to the reference sequence of*bin3* (NCBI Reference Sequence NM_165468.2). These 3 constructs were recombined into the *attP* landing site on chromosome 3L in BDSC stock #8622 (*y^1^ w^67c23^; P{y^+t7.7^=CaryP}attP2*) by BestGene, Inc.

### Expression of *UAS-DSCP-3xHA-bin3* transgenes using the Trojan GAL4 approach

We used the UAS/GAL4 system (Brand and Perrimon 1993) and the Trojan GAL4 approach (Diao and White 2012; Diao *et al*. 2015; Lee *et al*. 2018) to induce the expression of *UAS-DSCP-3xHA-bin3* (hereon referred to as *UAS-bin3*) transgenes. Intronic Trojan GAL4 insertions produce a null allele of a gene while also expressing GAL4 under the regulatory sequences of that gene. A Trojan GAL4 insertion into one of the *bin3* introns (*bin3^MI08045-TG4.0^*, hereon referred to as *bin3^TG4.0^*) in *trans* to a deficiency (*Df*) uncovering *bin3* produces *bin3*-null (*bin3^TG4.0^/Df*) flies that express GAL4 in the spatiotemporal pattern of endogenous *bin3*. (**Fig. 4a**). When transgenes comprising *UAS-bin3* cDNAs (**Fig. 5d-f**) are incorporated into *bin3^TG4.0^/Df* flies, GAL4 induces expression of Bin3 proteins in the pattern of endogenous *bin3* in an otherwise *bin3*-null background.

### Total RNA purification from ovaries and RT-qPCR

Five one to three-day old females were incubated with males in yeasted vials overnight. Five pairs of ovaries were dissected from two to four-day old females in ice-cold 1X PBS, and flash-frozen in liquid nitrogen. Total RNA was extracted using the Monarch Total RNA Miniprep Kit (NEB), according to the manufacturer’s instructions for RNA purification from “Tissues”. RT-qPCR was performed on 10 ng total RNA using the Luna Universal One-Step RT-qPCR Kit (NEB, E3005L). *bin3*, 7SK, U6, and *rp49* mRNAs were reverse transcribed and amplified with primers UP0493 + UP0118, UP0146 + UP0147, OW1123 + OW1124, and UP0113 + UP0114, respectively (see**Table S4** for primer sequences). Reactions were performed using a CFX Opus 384 Real-Time PCR System (Bio-Rad).

### Protein extraction and Western Analysis

Ten one to three-day old females were incubated with males in yeasted vials overnight. Ten pairs of ovaries were dissected from two to four-day old females in ice-cold 1X PBS, and flash-frozen in liquid nitrogen. Protein was exacted as previously described {Prudêncio.2015}. Extracts were clarified through 0.22 µM Spin-X columns (Costar), and mixed 1:1 with 2X Sample Loading Buffer (Bio-Rad) + 5% β-mercaptoethanol, and boiled for 5 minutes. Extracts were spun down, and 15 µl was loaded onto 4-20% TGX gels (Bio-Rad), and ran at 100 V for 10 minutes, then 300 V for 20 minutes. Protein was transferred to PVDF-LF membranes (Bio-Rad) using a Trans-Blot Turbo Transfer System (Bio-Rad) using the High MW preset (1.3 A, 25 V, 10 minutes). Membranes were blocked in EveryBlot Blocking Buffer (Bio-Rad) for 5 minutes, and then incubated with primary antibodies diluted in EveryBlot Blocking Buffer for 1 hour at room temperature. Membranes were washed with TBST (1X Tris-Buffered Saline, 0.2% Tween-20) for 5 minutes 3 times, and then incubated with secondary antibodies diluted in EveryBlot Blocking Buffer for 1 hour at room temperature. Membranes were with TBST for 5 minutes 3 times, dehydrated with 100% methanol, and imaged using a ChemiDoc MP Imaging System with Image Lab Touch software (Bio-Rad). Primary antibodies used were 1 µg/ml rabbit anti-HA (abcam, ab9110), and 1 µg/ml mouse anti-beta Actin (abcam, ab170325), and 5 µg/ml IgG purified from sheep anti-CycT antiserum (Nguyen *et al*. 2012). using Pierce Protein G Magnetic Beads according to the manufacturer’s instructions. Secondary antibodies were from Jackson ImmunoResearch, Inc., and used at 1 µg/ml: Alexa Fluor 488 AffiniPure Donkey Anti-Rabbit IgG (711-545-152), Alexa Fluor 488 AffiniPure Donkey Anti-Sheep IgG (713-545-147), Alexa Fluor 594 AffiniPure Donkey Anti-Mouse IgG (715-585-150), and Alexa Fluor 647 AffiniPure Donkey Anti-Mouse IgG (715-605-150).

### Egg-Laying Assay

Ten one to three-day old virgin females were incubated with wild-type males *en masse* in yeasted vials overnight. Ten two to four-day old females were split individually into fresh vials with two Oregon-R males, and incubated for 24 hours. After 24 hours, eggs were counted to determine the number of eggs laid/female/day. Experiments were performed in biological triplicate.

### Immunofluorescence staining of ovaries

Immunofluorescence staining of ovaries was performed as described (Maimon and Gilboa 2011). Ten 1-4 day old females were incubated with males in yeasted vials overnight, and then 10 pairs of ovaries were dissected from 2-5 day old females in ice-cold 1X PBS. The following steps were performed at RT unless otherwise indicated. Ovaries were fixed in 1 ml of Image-iT Fixative Solution (Thermo Scientific, R37814) + 0.3% Triton X-100 for 20 minutes. Ovaries were washed and permeabilized with 1 ml of 1% PT Buffer (1X PBS, 1% Triton X-100) for 5 minutes, 10 minutes, and 45 minutes. Ovaries were blocked with 1 ml PBT (1X PBS, 1% BSA, 0.3% Triton X-100) for 1 hour, and then incubated in 250 µl PBT and primary antibodies overnight at 4°C. Primary antibodies used were 1 µg/ml rabbit anti-HA (abcam ab9110) and 5 µg/ml mouse anti-Vasa (Developmental Studies Hybridoma Bank, Vasa 46F11). Ovaries were washed with 1 ml 0.3% PT Buffer (1X PBS, 0.3% Triton X-100) 3 times for 30 minutes each, blocked with 1 ml PBT-NDS (1X PBS, 1% BSA, 0.3% Triton X-100, 5% normal donkey serum) for 1 hour, and incubated in 250 µl PBT-NDS containing 10 µg/ml Hoechst 33342 (Thermo Scientific, H3570) and 2 µg/ml secondary antibodies for 2 hour in the dark. Secondary antibodies were from Jackson ImmunoResearch, Inc.: Alexa Fluor 488 AffiniPure Donkey Anti-Rabbit IgG (711-545-152) and Alexa Fluor 647 AffiniPure Donkey Anti-Mouse IgM (715-605-020). Ovaries were washed with 1 ml 0.3% PT Buffer in the dark 3 times for 30 minutes each, then mounted in 50 µl VECTASHIELD PLUS Antifade Mounting Medium (Vector Labs, Inc., H-1900-10).

### Single-molecule fluorescence *in situ* hybridization (smFISH) in ovaries

smFISH on adult ovaries was performed essentially as described (Raj and Tyagi 2010; Abbaszadeh and Gavis 2016). Ten 1-4 day old females were incubated with males in yeasted vials overnight, and then 10 pairs of ovaries were dissected from 2-5 day old females in ice-cold 1X PBS. The following steps were performed at RT unless otherwise indicated. Ovaries were fixed in 1 ml of Image-iT Fixative Solution (Thermo Scientific, R37814) for 30 minutes, then washed with PBST (1X PBS, 0.1% Tween-20) 3 times for 5 minutes each. Ovaries were permeabilized by incubating in 7:3 PBST:methanol for 5 minutes, 1:1 PBST:methanol for 5 minutes, 3:7 PBST:methanol for 5 minutes, 100% methanol for 10 minutes, 3:7 PBST:methanol for 5 minutes, 1:1 PBST:methanol for 5 minutes, and 7:3 PBST:methanol for 5 minutes. Ovaries were then washed with PBST 4 times for 5 minutes each, followed by pre-hybridized in Wash Buffer A (20% Stellaris RNA FISH Wash Buffer A (Biosearch Technologies Cat# SMF-WA1-60) and 10% deionized formamide) for 5 minutes. Ovaries were incubated in Stellaris RNA FISH Hybridization Buffer (Biosearch Technologies Cat# SMF-HB1-10) with 10% deionized formamide and 125 µM Quasar 640-labeled oligonucleotide probes, overnight at 37°C in the dark. Oligonucleotide probes against 7SK were designed using the LGC Biosearch Technologies’ Stellaris RNA FISH Probe Designer (https://www.biosearchtech.com/support/tools/design-software/stellaris-probe-designer). Ovaries were washed in the dark with Wash Buffer A for 30 minutes at 37°C, and then with Wash Buffer A with 10 µg/ml Hoechst 33342 (Thermo Scientific, H3570) for 30 minutes at 37°C, and finally with Stellaris RNA FISH Wash Buffer B (Biosearch Technologies Cat# SMF-WB1-20) for 5 minutes at RT. Ovaries were mounted in 50 µl VECTASHIELD PLUS Antifade Mounting Medium (Vector Labs, Inc., H-1900-10).

### Ovariole Counting Assay

One to three-day old females were incubated with sibling males in yeast vials overnight. Two to four-day females were anesthetized, transferred to 1.7 ml tubes, flash-frozen in liquid nitrogen, and stored at −80°C. Frozen females were thawed on ice or at RT and dissected in a 1:1 mixture of methanol and 1X PBS. 50% methanol dehydrates and extracts lipids from the gonadal sheath surrounding the ovary and from the clear portions of the ovarioles (turning them opaque), making ovarioles easy to separate and count. The mean number of ovarioles per ovary, as well as the absolute value of the difference in ovariole number between each ovary per female were calculated (Lobell *et al*. 2017). Experiments were performed in biological duplicate.

### Climbing Assay

Ten 2-5 day old females were transferred to 15 ml tubes, and allowed to acclimate for 3’. Flies were tapped to the bottom of the tube, and climbing was recorded for 12 seconds. Flies were then allowed to reacclimatize for another 3 minutes before performing the subsequent trial. 10 trials were performed per genotype to determine the percent of flies climbing 8 cm in 12 seconds per trial. All genotypes for a particular experiment were tested at the same time. Experiments were performed in biological triplicate.

### Modeling of Bin3 protein sequence onto MEPCE

Bin3 was modeled onto 309 amino acids of the MEPCE active site in complex with *S* -adenosyl-homocysteine (SAH) and stem loop 1 of 7SK (PDB 6dcb; Yang *et al*. 2019) using hhpred (Söding 2005; Hildebrand *et al*. 2009; Meier and Söding 2015; Zimmermann *et al*. 2018; Gabler *et al*. 2020) and MODELLER (Šali *et al*. 1995; Zimmermann *et al*. 2018; Gabler *et al*. 2020). Alignment of the Bin3 model to 6dcb, RMSD calculation, and modeling of the Y421A and Y795A mutations was performed using PyMol (version 2.5.4).

### Bin3 Disorder Prediction

Disorder in Bin3 was predicted by IUPred3, using the IUPred3 short disorder analysis type (Mészáros *et al*. 2018; Erdős and Dosztányi 2020; Erdős *et al*. 2021).

### RNA Immunoprecipitation (RIP)

Twenty-five one to three-day old females with or without transgenes expressing *bin3-3xHA* cDNAs were incubated with males in yeasted vials overnight. Twenty-five pairs of ovaries were dissected from two to four-day old females in ice-cold, 1X PBS, and fixed in 1 ml 1.8% formaldehyde (a mixture of 55% 1X PBS and 45% Image-iT Fixative Solution, Thermo Scientific, R37814) for 15 minutes at RT on a nutating rotor. Fixation was terminated by addition of 163 µl 2.5 M glycine to a final concentration of ∼350 mM, and incubation for 5 minutes at RT on a nutating rotor. Ovaries were washed briefly with with 1X PBS 3 TIMES, and then flash-frozen in liquid nitrogen. Ovaries were homogenized by hand with a blue pestle on ice in 50 µl ice-cold RIPA Lysis and Extraction Buffer (Thermo Scientific, 89900) supplemented with 1 mM DTT, 3X Halt Protease and Phosphatase Inhibitor Cocktail (Thermo Scientific, 78440), and 1 U/µl SUPERase•In RNase Inhibitor (Invitrogen, AM2696). 200 µl of additional buffer was added to the homogenate, and samples were incubated on ice for 15 minutes to allow for complete nuclear lysis. Homogenate was pelleted by centrifugation at top speed for 1 minute. 2.5 µl of homogenate (1/100^th^ homogenate volume) was removed and kept on ice in a total of 300 µl 1X Monarch DNA/RNA Protection Reagent (NEB, T2011L) as the input sample. The remaining homogenate was used for immunoprecipitation. 25 µl (1/10^th^ homogenate volume) of Pierce Anti-HA Magnetic Beads (Thermo Scientific, 88836) were washed with 175 µl 1X TBS-T (Pierce 20X TBS Tween 20 Buffer, Thermo Scientific, 28360), and then with 1 ml 1X TBS-T. Homogenate was added to the washed beads, and immunoprecipitation was performed for 4 hours at 4°C with end-over-end rotation. Beads were washed briefly with 1 ml 1X TBS-T 3 times. Bin3-3xHA complexes were purified by two sequential competitive elutions with 75 µl of Pierce HA Peptide (Thermo Scientific, 26184) diluted to 2 mg/ml in Nuclease-Free Water (NEB, B1500L), at 37°C for 15 minutes each. The eluates were pooled, and the 150 µl total eluate was added to 150 µl 2X Monarch DNA/RNA Protection Reagent. All samples (inputs and eluates) were incubated at 65-70°C for 30 minutes to reverse formaldehyde crosslinks. Total RNA was extracted using the Monarch Total RNA Miniprep Kit (NEB), according to the manufacturer’s instructions for RNA purification from “Tissues”. Total RNA was eluted in 100 µl Nuclease-Free Water. 2.9 µl of each sample was used in 10-µl reactions to reverse transcribe and amplify 7SK and U6 snRNAs with primers UP0146 + UP0147 and OW1123 + OW1124, respectively (see**Table S4** for primer sequences). Reactions were performed using the Luna Universal One-Step RT-qPCR Kit (NEB, E3005L) and a CFX Opus 384 Real-Time PCR System (Bio-Rad).

## RESULTS

### Bin3 is required for female fecundity

Reproductive success is influenced by fertility (offspring viability) and fecundity (the rate of offspring production). Previously, we showed that Bin3 plays a role in fertility, promoting embryonic viability by a mechanism of *caudal* translation repression (Singh *et al*. 2011). During that and other studies, we noticed that *bin3* mutants lay fewer eggs than wild type flies. Therefore, we asked if Bin3 also plays a role in fecundity by promoting a normal egg-laying rate. To do this, we compared the egg-laying rates of flies that were either wild type for *bin3* (wt), heterozygous for *bin3* (*bin3^2-7^/+*), or deleted entirely for *bin3* (*bin3^2-7^* in *trans* to a deficiency that removes the *bin3* locus; *bin3^2-7^/Df*). The genomic regions deleted by *bin3^2-7^* and *Df* are depicted in **Fig. 1a**. As expected, *bin3* mRNA levels are reduced by half in *bin3^2-7^/+* heterozygotes and are undetectable in *bin3^2-7^/Df* mutants (**Fig. 1b**). Interestingly, we found that *bin3^2-7^/Df* females laid eggs at a significantly reduced rate compared to wt (*P* =.0001) and *bin3^2-7^/+* (*P* <.0001) females (**Fig. 2a**). The egg-laying rate defect was specific to the loss of *bin3*, as a single copy of a genomic fragment containing the *bin3* ORF (*bin3^+^*, **Fig. 1a**) restored *bin3* mRNA accumulation by half (**Fig. 1b**) and rescued the egg-laying defect of *bin3^2-7^/Df* females (*P* <.0001, **Fig. 2a**). Rescue was comparable to that of *bin3^2-7^/+* controls. These results demonstrate that Bin3 is required for egg-laying.

**Figure 2.**
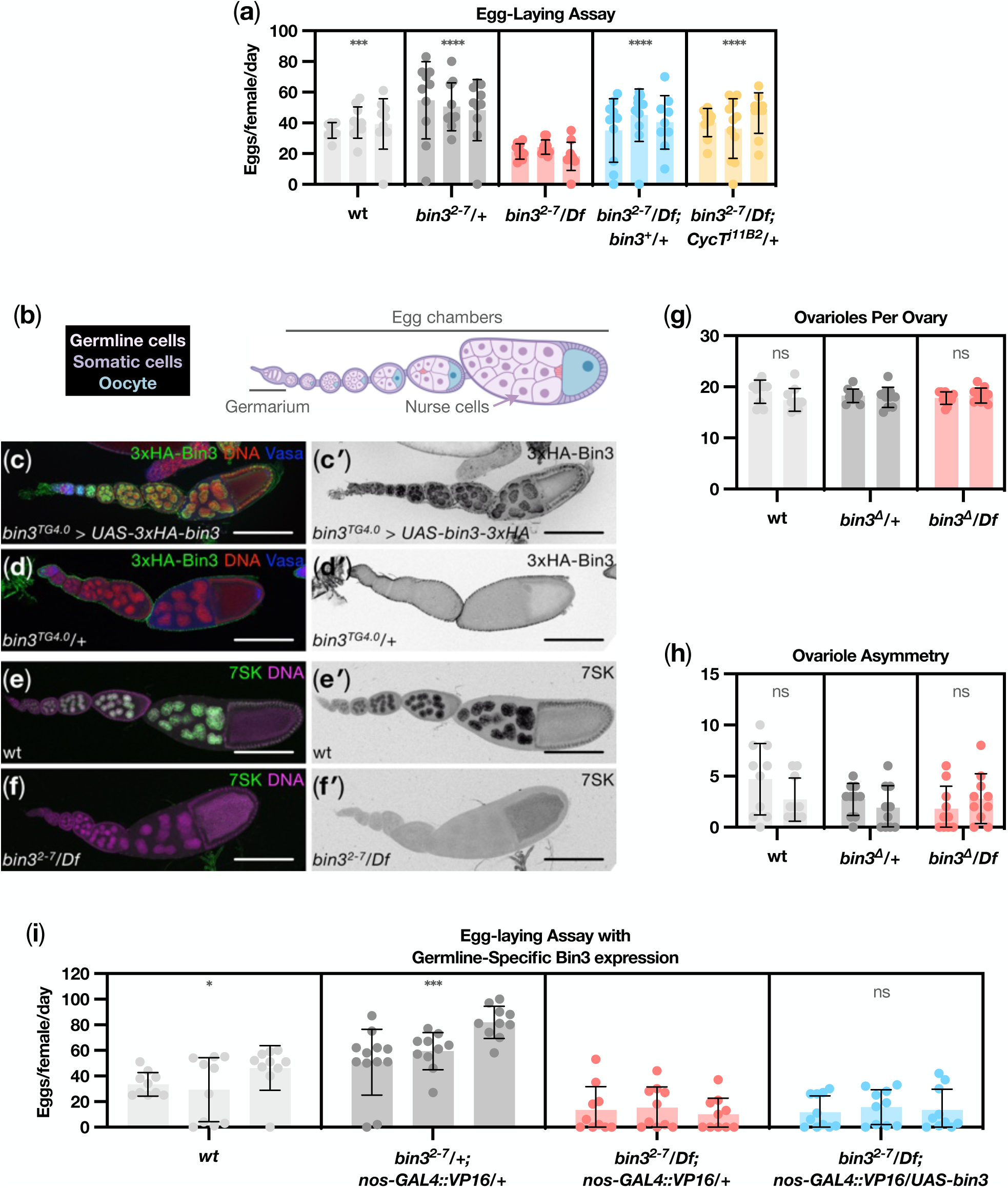
Bin3 is ubiquitously expressed in the ovary and is required for egg-laying, but is not required to establish ovarioles and does not function in germline to promote egg-laying. **(a)** Scatter plot showing the egg-laying rates of females of the indicated genotypes for each of three biological replicates. Error bars represent the standard deviation. Statistical significance compared to *bin3^2-7^/Df* was calculated by nested one-way ANOVA with Dunnett ’s Correction for multiple comparisons, using Prism (version 9.4.1, GraphPad). *P* values of <.001 (***) and <.0001 (****) are indicated. **(b)** Cartoon depicting a single ovariole, of which 16-20 make up a single adult ovary. Germline cells are indicated in pink, nurse cell nuclei (arrow) are indicated in purple, oocytes are indicated in blue, and somatic cells surrounding the germline and oocytes are indicated in purple. **(c to d’)** Immunofluorescence staining of adult ovaries for 3xHA-Bin3 (green), Vasa (a cytoplasmic marker for germline cells; blue), and Hoechst staining of DNA (red). Scale bars = 100 µm. A Trojan-GAL4 phase 0 insertion in *bin3* (*bin3^TG4.0^*) was used to drive expression of a *UAS-3xHA-bin3* transgene in the pattern of *bin3* expression (*bin3^TG4.0^ > UAS-bin3-3xHA*). 3xHA-Bin3 localizes to the nuclei of all germline and somatic cells (C-C’). No HA staining was detected in *bin3^TG4.0^*control ovaries (D-D’), demonstrating the specificity of HA staining in *bin3^TG4.0^ > UAS-3xHA* -*bin3* ovaries. **(e-f’)** Single-molecule fluorescence *in situ* hybridization (smFISH) of adult ovaries for 7SK (green) and Hoechst staining of DNA (purple). Scale bars = 100 µm. 7SK localizes to the nuclei of all germline and somatic cells in wt ovaries (E-E’). 7SK staining is absent in *bin3^2-7^/Df* ovaries (F-F’), which do not accumulate 7SK (see Fig. 1c). **(g)** Scatter plot showing the mean number of ovarioles per ovary per female of the indicated genotypes for each of two biological replicates. Error bars represent the standard deviation. Statistical significance compared to *bin3^2-7^/Df* was calculated by nested one-way ANOVA with Dunnett ’s Correction for multiple comparisons, using Prism (version 9.4.1, GraphPad). *P* values of not significant (ns) are indicated. **(h)** Scatter plot showing the absolute value of the difference between ovariole number within a pair of ovaries per female of the indicated genotypes for each of two biological replicates. Error bars represent the standard deviation. Statistical significance compared to *bin3^2-7^/Df* was calculated by nested one-way ANOVA with Dunnett ’s Correction for multiple comparisons, using Prism (version 9.4.1, GraphPad). *P* values of not significant (ns) are indicated. **(i)** Scatter plot showing the egg-laying rates of females of the indicated genotypes for each of three biological replicates. Error bars represent the standard deviation. Statistical significance compared to *bin3^2-7^/Df; nanos-GAL4::VP16/+* was calculated by nested one-way ANOVA with Dunnett’s Correction for multiple comparisons, using Prism (version 9.4.1, GraphPad). *P* values of <.05 (*), <.001 (***) and not significant (ns) are indicated.

### Bin3 promotes female fecundity by repressing P-TEFb

We found that 7SK RNA levels were strongly reduced in *bin3^2-7^/Df* females compared to wt (*P* <.0001, **Fig. 1c**), as we have previously shown (Singh *et al*. 2011). The canonical function of Bin3/MEPCE is to stabilize 7SK RNA (Jeronimo *et al*. 2007; Singh *et al*. 2011). 7SK RNA in turn acts as a scaffold for proteins that form a snRNP that sequesters and represses P-TEFb (Krueger *et al*. 2008; Barboric *et al*. 2009; Xue *et al*. 2010; Muniz *et al*. 2013; Brogie and Price 2017; Yang *et al*. 2022). Therefore, destabilization of 7SK in *bin3^2-7^/Df* females should lead to deregulation (activation) of P-TEFb, which could be responsible for the egg-laying defect. If this is true, then we expect that reducing the activity of P-TEFb in *bin3^2-7^/Df* females would rescue the egg-laying defect. To genetically reduce the activity of P-TEFb, we introduced a mutant allele of *CycT* (*CycT^jllB2^*) into *bin3^2-7^/Df* flies, and confirmed that this mutation reduces the protein levels of the largest of the two CycT isoforms (**Fig. 1e**). Indeed, we found that reducing the amount of CycT in *bin3^2-7^/Df* females rescued the egg-laying defect (P < .0001, **Fig. 2a**), while having no effect on 7SK RNA levels (**Fig. 1c**). These results suggest that Bin3 stabilizes 7SK RNA to form the 7SK snRNP, which then sequesters and represses P-TEFb to promote a normal egg-laying rate.

### Bin3 and 7SK are expressed ubiquitously in the ovary

To gain insight into the role of Bin3 in egg-laying, we determined where Bin3 (and its substrate 7SK) is expressed and localized in ovaries, which are composed of 16-20 individual ovarioles that can be thought of as assembly lines of germline development (**Fig. 2b**). We used the UAS/GAL4 (Brand and Perrimon 1993) and Trojan GAL4 systems (see Materials and Methods; Diao and White 2012; Diao *et al*. 2015; Lee *et al*. 2018), where a Trojan GAL4 insertion into the *bin3* ORF was used to drive the expression of a 3xHA-epitope-tagged *bin3* transgene in the spatiotemporal pattern of endogenous *bin3* (*bin3^TG4.0^ > UAS-3xHA-bin3*). Bin3-3xHA was detected by immunofluorescence staining of ovaries. We found that Bin3 is expressed in and localized to the nucleic of all ovarian cells (**Fig. 2, c, c’**). Bin3 was found to be expressed in the germline cells of the germarium (located at the anterior of each ovariole), and in the 15 nurse cells that supply the oocyte with mRNAs and proteins. Bin3 was also found to be expressed in the somatic cells surrounding the germline cells and the oocyte, which communicate with germline cells to promote egg chamber and oocyte maturation. Control flies lacking the *UAS-3xHA-bin3* transgene (*bin3^TG4.0^/+*) yielded no detectable HA staining (**Fig. 2, d, d’**).

To identify where in the ovary 7SK RNA is expressed and localized, we performed single-molecule fluorescence *in situ* hybridization (Raj and Tyagi 2010; Abbaszadeh and Gavis 2016) on ovaries from wt and *bin3^2-7^/Df* females. We found that in wt ovarioles, 7SK follows an identical expression pattern to that of Bin3 (compare **Fig. 2, e, e’** with **Fig. 2, c, c’**). As expected, 7SK staining was undetectable in *bin3^2-7^/Df* ovarioles (**Fig. 2, f, f’**), which do not accumulate 7SK RNA (**Fig. 1c**). Taken together, Bin3 and 7SK are both expressed in all cells of the ovarioles and localize to nuclei— commensurate with their role in transcription elongation regulation.

### Bin3 is not required to establish the proper number of ovarioles

The total number of ovarioles per ovary directly correlates with the number of eggs insects can lay (Wayne *et al*. 1997; Wayne and Mackay 1998), and is established by proper proliferation and migration of the somatic terminal filament cells (TFCs; Sahut-Barnola *et al*. 1995, 1996). Given that we found *bin3^2-7^/Df* mutant females had an egg-laying defect, and that Bin3 is expressed in the somatic cells of the ovarioles, we hypothesized that Bin3 promotes egg-laying by functioning in the TFCs to establish ovarioles. To test this hypothesis, we used the UAS/GAL4 system (Brand and Perrimon 1993) with two different *bab1-GAL4* drivers (Cabrera *et al*. 2002) to induce expression of the *UAS-3xHA-bin3* transgene specifically in the somatic TFCs of *bin3^2-7^/Df* females to determine whether this rescued the egg-laying rate. Unfortunately, we were unable to recover *bin3^2-7^/Df; bab1-GAL4/UAS-bin3* flies using either driver. Both *bab1-GAL4* drivers are expressed in several cell types in addition to the TFCs (Cabrera *et al*. 2002), and therefore it is likely that ectopic expression of *bin3* in all *bab1*-expressing cells is lethal.

As an alternative approach, we compared ovariole numbers in *bin3^2-7^/Df* females with those of wt and *bin3^2-7^/+* controls. To do this, we dissected ovaries and determined (i) the average number of ovarioles per ovary, and (ii) the difference in the number of ovarioles within a pair of ovaries (i.e., “ovariole asymmetry”; Lobell *et al*. 2017). Unexpectedly, we did not find a significant difference in either the mean number of ovarioles per ovary (**Fig. 2g**) or in ovariole asymmetry (**Fig. 2h**) in *bin3^2-7^/Df* females compared to either wt or *bin3^2-7^/+* control females.

Together, these results suggest that Bin3 is not required to establish ovarioles, and therefore promotes egg-laying by some other mechanism. These results also suggest that Bin3 does not have a function in TFCs relevant to the establishment of ovarioles. However, these results do not rule out some other function of Bin3 in TFCs (*e.g*., cell-cell communication with germline cells) that might promote egg-laying. The lethality of inducing *bin3* expression in TFCs using the *bab1-GAL4* drivers precludes the ability to test this hypothesis in any straightforward way.

### Expression of Bin3 in the germline is not sufficient to promote fecundity

Bin3 is also expressed in germline cells, therefore, it remained possible that Bin3 functions in the germline to promote a normal egg-laying rate. To test this hypothesis, we used a *nanos-GAL4::VP16* driver (Doren *et al*. 1998) to induce expression of the *UAS-3xHA-bin3* transgene specifically in the germline cells of *bin3^2-7^/Df* females, and asked whether this improved the egg-laying rate (**Fig. 2g**). Results showed that germline expression of *bin3* did not rescue the egg-laying rate defect of *bin3^2-7^/Df; nanos-GAL4::VP16/UAS-bin3* females compared to *bin3^2-7^/Df; nanos-GAL4::VP16/+* mutant females (*P* = .9942, **Fig. 2i**), suggesting that Bin3 does not function in germline cells to promote a normal egg-laying rate.

### Bin3 has a conserved role in neuromuscular function

A nonsense mutation in the *MEPCE* gene of a patient with impaired neuromuscular ability was shown to reduce MEPCE protein levels, causing P-TEFb hyper-activation and aberrant RNAPII transcription elongation (Schneeberger *et al*. 2019). If a conserved function of Bin3/MEPCE is to repress P-TEFb to promote normal neurodevelopment, then it would be expected that *bin3* mutant flies would also have neuromuscular defects, and that these defects should be rescued by reducing CycT levels (lowering P-TEFb activity).

Two ways in which neuromuscular function can be assessed in flies is by examining climbing (Manjila and Hasan 2018) and resting wing posture (Baehrecke 1997). Normally, flies climb to the top of an enclosure after being knocked to the bottom, and fold their wings one-over-the-other when resting. We found that a significant percentage of *bin3^2-7^/Df* flies had a climbing defect compared to *bin3^2-7^/+* control flies (P = .0003, **Fig. 3a**). Additionally, whereas control *bin3^2-7^/+* flies properly folded their wings (**Fig. 3b**), *bin3^2-7^/Df* mutant flies had a *held-out wings* phenotype (**Fig. 3c**). Both defects were specific to loss of *bin3*, as the *bin3^+^* genomic fragment rescued the climbing (*P* = .0004, **Fig. 3a**) and resting wing posture defects of *bin3^2-7^/Df* flies (compare **Fig. 3d** with **Fig. 3c**). Importantly, both climbing (*P* = .0008, **Fig. 3a**) and wing posture (compare **Fig. 3e** with **Fig. 3c**) defects of *bin3^2-7^/Df* flies were also rescued by reducing CycT levels. These results suggest that Bin3 and MEPCE share a conserved role in neuromuscular function from *Drosophila* to humans, most likely by repressing P-TEFb and preventing aberrant transcription elongation.

**Figure 3.**
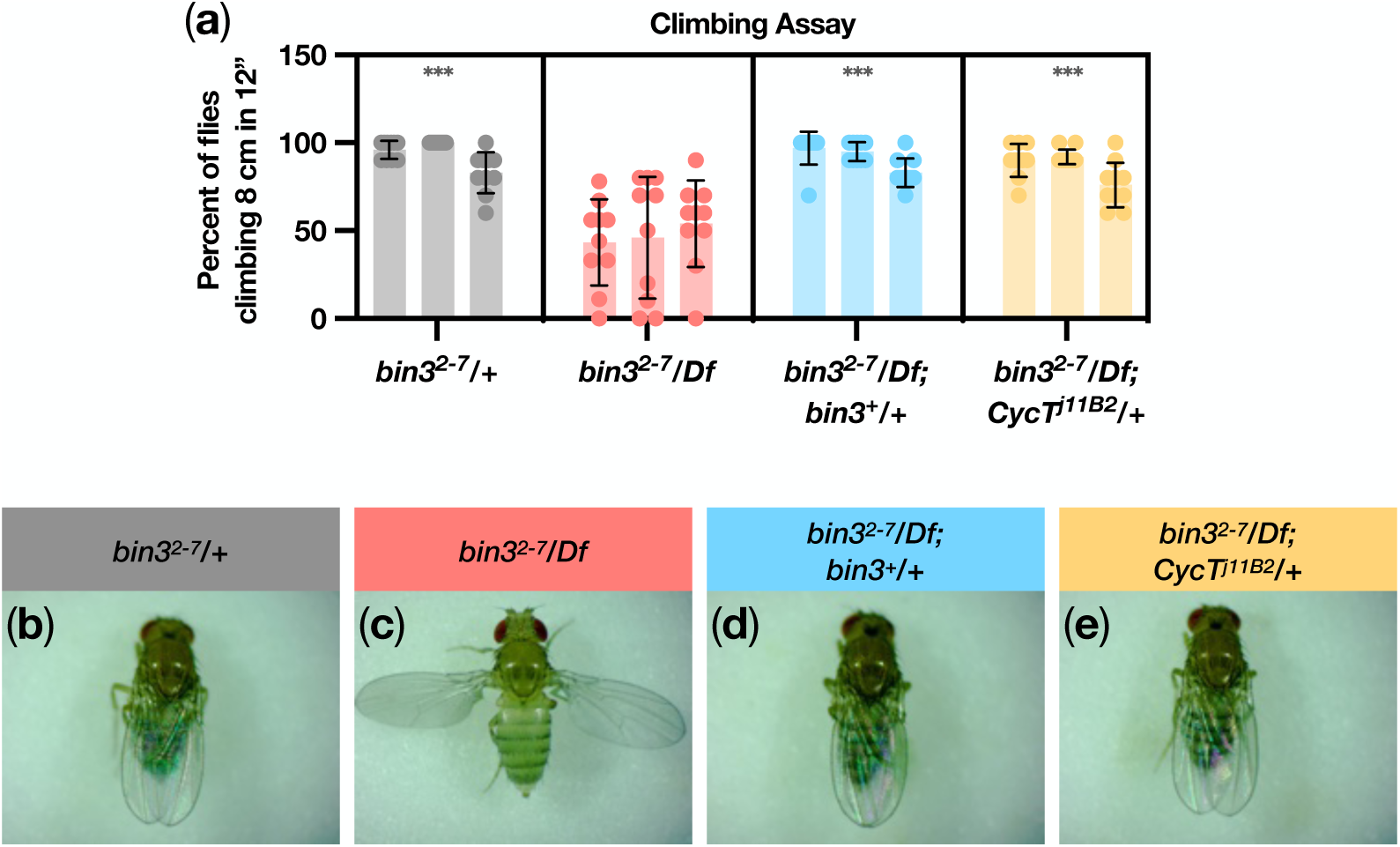
Bin3 is required for neuromuscular function. **(a)** Scatter plot showing the percent of flies of the indicated genotypes climbing 8 cm in 12” in each of 10 trials, for each of three biological replicates. Error bars represent the standard deviation. Statistical significance compared to *bin3^2-7^/Df* was calculated by nested one-way ANOVA with Dunnett ’s Correction for multiple comparisons, using Prism (version 9.4.1, GraphPad). *P* value of <.001 (***) is indicated. **(b-e)** Images of females of the indicated genotypes, exhibiting either a normal resting wing posture or a *held-out wings* phenotype. **(f)** Scatter plot showing the percent of flies of the indicated genotypes climbing 8 cm in 12” in each of 4 trials, for each of three biological replicates. Error bars represent the standard deviation. Statistical significance was calculated by nested t-test, using Prism (version 9.4.1, GraphPad). *P* value of not significant (ns) is indicated. **(g-h)** Images of females of the indicated genotypes, exhibiting either a normal resting wing posture or a *held-out wings* phenotype.

### Bin3 catalytic activity is dispensable for 7SK stability and snRNP function *in vivo*

The importance of MEPCE catalytic residues in 7SK binding and capping have been studied mostly *in vitro* (Xue *et al*. 2010; Shelton *et al*. 2018; Yang *et al*. 2019). To determine the importance of Bin3 catalytic activity *in vivo,* we sought to generate flies expressing a catalytically-dead Bin3 protein. We first analyzed the crystal structure of the methyltransferase domain of human MEPCE (PDB 6dcb, Yang *et al*. 2019). As shown in **Fig. 4a**, Tyrosine 421 (Y421) is juxtaposed between the 5’ gamma phosphate of 7SK and S-adenosyl-homocysteine (SAH; used in place of the S-adenosyl-methionine (SAM) methyl donor for the purposes of crystallization). Y421 hydrogen bonds to the 7SK gamma phosphate. Mutation of this tyrosine to alanine (Y421A) completely abolished MEPCE catalytic activity *in vitro* (Yang *et al*. 2019). We modeled Bin3 onto the crystal structure (**Fig. 4a**) and found that Y795 in Bin3 (red) aligned well with Y421 in MEPCE (black). We created models (**Fig. 4b**) representing the Y421A mutation in MEPCE (black) and the concomitant Y795A mutation in Bin3 (red), which revealed a complete loss of coordination between 7SK, SAH (i.e., SAM), and MEPCE/Bin3.

Given the *in vitro* importance of the Y421 residue in MEPCE for catalytic activity, and that Y795 in Bin3 aligns with Y421, we generated flies expressing either wt Bin3 (Bin3 ^wt^) or Bin3^Y795A^ and tested whether Bin3^Y795A^ was able to (i) bind and stabilize 7SK, and (ii) rescue *bin3* mutant phenotypes. To do this, we again used the Trojan GAL4 approach. Importantly, not only do Trojan GAL4 insertions express GAL4 in the pattern of the gene they are inserted into, but they simultaneously generate a null allele of the inserted gene. Indeed, *bin3* Trojan GAL4 insertion flies (*bin3^TG4.0^/Df*) (**Fig. 4d**) had nearly undetectable levels of both *bin3* mRNA (*P* < .0001, **Fig. 4e**) and 7SK RNA (*P* < .0001 **Fig. 4f**) compared to wt, confirming that *bin3^TG4.0^*is a null allele of *bin3*. Additionally, we found that GAL4 expressed from *bin3^TG4.0^* induced both *UAS-bin3^wt^* (**Fig. h**) and *UAS-bin3^Y795A^* (**Fig. 4i**) transgenes similarly at the mRNA (*P* = .2188, **Fig. 4k**) and protein (*P* = .9836, **Fig. 4l**) levels. Thus, these flies could be used to study the importance of Bin3 catalytic activity *in vivo*.

Bin3 catalytic activity is expected to be required for capping and stabilization of 7SK *in vivo*; therefore, in Bin3^Y795A^ mutant flies, 7SK should be destabilized. Remarkably, there was no significant difference in 7SK levels between Bin3^Y795A^ flies (*bin3^TG4.0^/Df; UAS-bin3^Y795A^/+*) and Bin3^wt^ flies (*bin3^TG4.0^/Df; UAS-bin3^wt^/+*; *P* = .9996, **Fig. 5a**), suggesting that Bin3 catalytic activity is not required for the stability of 7SK *in vivo*. How then might Bin3 stabilize 7SK in the absence of catalytic activity? As suggested by prior studies (Xue *et al*. 2010), Bin3 might form a stable complex with 7SK independent of its catalytic function, and this may protect it from degradation. Indeed, we find that Bin3 ^Y795A^ immunoprecipitated as much 7SK RNA as did Bin3 ^wt^ (*P* = .5350, **Fig. 5b**), suggesting that the Y795A mutation does not diminish the binding of Bin3 to 7SK, and that binding alone is sufficient for stabilization *in vivo*.

Given that Bin3 ^Y795A^ could bind and stabilize 7SK, we expected that Bin3 ^Y795A^ would rescue the fecundity, climbing, and wing posture defects of *bin3* mutant flies. First, we showed that *bin3^TG4.0^/Df* flies recapitulate the fecundity (compare **Fig. 6a** to **Fig. 2a**), climbing (compare **Fig. 6b** with **Fig. 3a**), and wing posture (compare **Fig. 6d** with **Fig. 3c**) defects of *bin3^2-7^/Df* flies. As expected, Bin3^Y795A^ expression rescued the fecundity (*P* = .0008, **Fig. 6a**), climbing (*P* < .0001, **Fig. 6b**), and wing posture (compare **Fig. 6f** to **Fig. 6d**) defects of *bin3^TG4.0^/Df* flies, similar to Bin3^wt^ (*P* > .9999 for both fecundity and climbing). Therefore, despite the importance of catalytic activity to 7SK capping *in vitro*, Bin3 catalytic activity and therefore the methylphosphate cap do not seem to be required *in vivo* for 7SK stability, nor is it likely to affect 7SK snRNP assembly and function (**Fig. 6i**).

**Figure 4.**
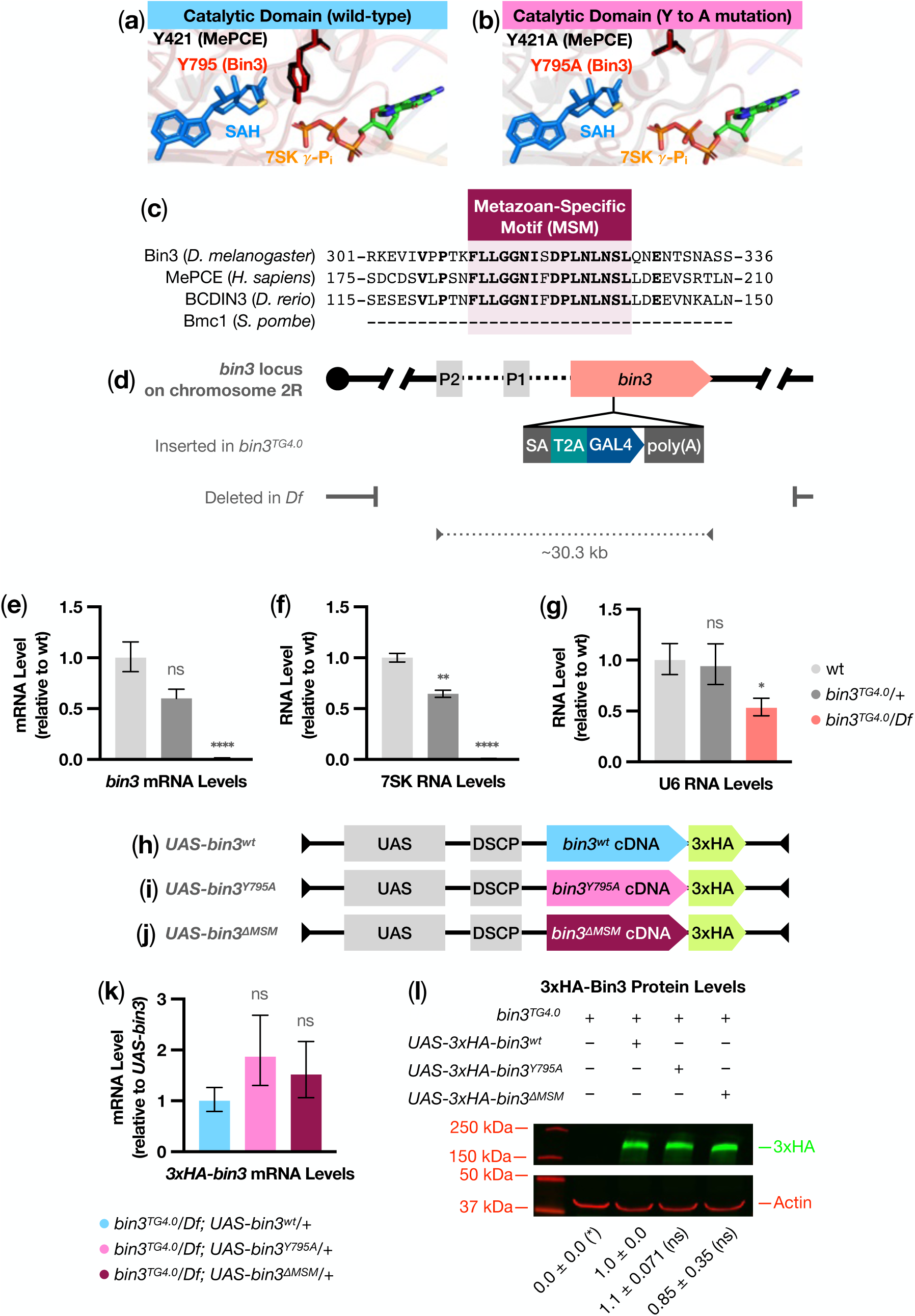
Molecular characterization of the *bin3^TG4.0^* Trojan GAL4 allele and Bin3 protein mutants. **(a)** Bin3 modeled onto the crystal structure of 309 amino acids of the MEPCE active site (Yang *et al*. 2019). Y421 in MEPCE (black) is juxtaposed between the 7SK 5’ gamma phosphate (orange and salmon) and *S* -adenosyl-homocysteine (SAH, blue and gold), and is essential for MEPCE catalytic activity *in vitro* (Yang *et al*. 2019). Y795 in Bin3 (red) aligns with Y421 in MEPCE (RMSD = 0.000). **(b)** Same as in (a), but with the Y421A (MEPCE, black) and Y795A (Bin3, red) tyrosine-to-alanine mutations introduced. Y421A completely eliminated MEPCE catalytic activity *in vitro* (Yang *et al*. 2019). **(c)** Partial alignment of Bin3 orthologs from *D. melanogaster*, *H. sapiens* (MEPCE), *D. rerio* (BCDIN3), and *S. pombe* (Bmc1). Conserved residues are in boldface. **(d)** Graphic depicting the *bin3* locus, as in Fig. 1a . *bin3^TG4.0^*refers to a transgenic insertion of a Trojan GAL4 phase 0 (TG4.0) construct into the second intron in the *bin3* ORF (Lee *et al*. 2018). A splice acceptor (SA) in the insertion causes the TG4.0 cassette to be spliced into the *bin3* mRNA; transcription is terminated at the end of the cassette by a synthetic poly(A) sequence. When the chimeric *bin3^TG4.0^* mRNA is translated, T2A, a viral peptide sequence, disrupts translation and causes the release of the nonfunctional Bin3:T2A chimeric peptide from the ribosome; subsequently, the ribosome resumes translation of full-length GAL4 protein. *Df* refers to deficiency *Df(2R)BSC313*, which deletes ∼625 kb of chromosome 2R, including promoters P2 and P1, and all *bin3* coding sequence (Cook *et al*. 2012). *bin3^TG4.0^* and *Df* in *trans* produce *bin3* mutant flies that express *GAL4* in the pattern of *bin3* expression. GAL4 then binds to UAS and drives expression of *UAS-bin3* transgenes (see Fig. 4h**-j**). **(e-g)** Bar graphs showing the average amount of *bin3* mRNA (e), 7SK snRNA (f), and U6 snRNA (g) in total RNA extracted from ovaries of females of the indicated genotypes, for three biological replicates. RNA levels were normalized to *rp49* mRNA and made relative to wt using CFX Maestro 2.0 (version 5.2.008.0222, Bio-Rad). Error bars represent the standard error of the mean. Statistical significance was calculated by unpaired t-test using CFX Maestro 2.0 (version 5.2.008.0222, Bio-Rad). *P* values of <.05 (*), <0.01 (**), <.0001 (****), and not significant (ns) are indicated. **(h-j)** Graphic depicting *UAS-DSCP-3xHA-bin3* transgenes. UAS-DSCP (grey) is a GAL4-responsive promoter that is active in both germline and somatic cells (Pfeiffer *et al*. 2008). *bin3* cDNAs encode Bin3 proteins that are either wt (h; sky blue), catalytically-dead (I; Y795A, pink), or deleted of the MSM (j; ΔMSM, maroon). Each construct is tagged at the C-terminus with 3xHA that is codon-optimized for *Drosophila* (green). When these transgenes are independently crossed into the *bin3^TG4.0^/Df* background (see Fig. 4d), flies will express only 3xHA-Bin3, 3xHA-Bin3 ^Y795A^, or 3xHA-Bin3^ΔMSM^ proteins, respectively. **(k)** Bar graph showing the average amount of *bin3* mRNA in total RNA extracted from ovaries of females of the indicated genotypes, for three biological replicates. RNA levels were normalized to *rp49* mRNA and made relative to *bin3^TG4.0^/Df; UAS-bin3/+*using CFX Maestro 2.0 (version 5.2.008.0222, Bio-Rad). Error bars represent the standard error of the mean. Statistical significance was calculated by unpaired t-test using CFX Maestro 2.0 software (version 5.2.008.0222, Bio-Rad). *P* value of not significant (ns) is indicated. All three *UAS-bin3* transgenes are expressed to similar levels. **(i)** Representative Western blot for 3xHA-Bin3 proteins and Actin in extracts prepared from ovaries of females of the indicated genotypes. The relative amounts of 3xHA-Bin3 ^Y795A^ and 3xHA-Bin3^ΔMSM^ relative to 3xHA-Bin3 (wt) ± the standard deviation from two biological replicates are indicated below. Statistical significance was calculated by ordinary one-way ANOVA with Dunnett ’s Correction for multiple comparisons with a single pooled variance, using Prism (version 9.4.1, GraphPad). *P* values of <0.05 (*) and not significant (ns) are indicated. All three transgenes produce similar amounts of 3xHA-Bin3 proteins.

**Figure 5.**
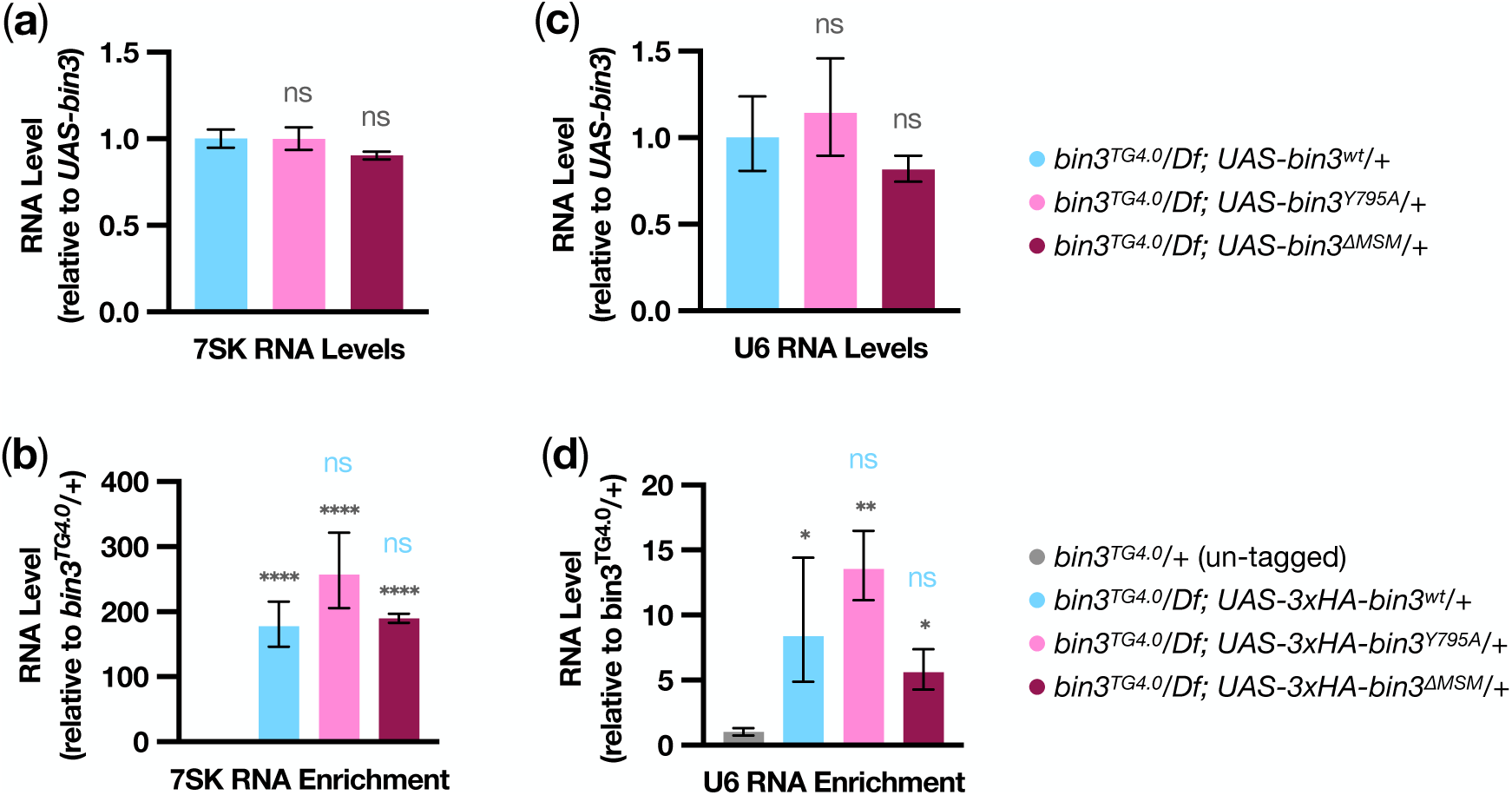
Neither catalytic activity nor the metazoan-specific motif are required for 7SK or U6 binding and stability. **(a** and **c)** Bar graphs showing the average amount of 7SK (a) and U6 (c) RNAs in total RNA extracted from ovaries of females of the indicated genotypes, for three biological replicates. RNA levels were normalized to *rp49* mRNA and made relative to *bin3^TG4.0^/Df; UAS-bin3/+* using CFX Maestro 2.0 (version 5.2.008.0222, Bio-Rad). Error bars represent the standard error of the mean. Statistical significance was calculated by unpaired t-test using CFX Maestro 2.0 (version 5.2.008.0222, Bio-Rad). *P* value of not significant (ns) is indicated. **(b** and **d)** Bar graphs showing the average amount of 7SK (b) and U6 (d) RNA bound by 3xHA-Bin3 proteins immunoprecipitated with anti-HA magnetic beads from extracts prepared from ovaries of females of the indicated genotypes, for three biological replicates. The ratio of immunoprecipitated RNA to input RNA, from flies expressing 3xHA-Bin3 proteins relative to flies not expressing 3xHA-Bin3 proteins (*bin3^TG4.0^/+*), was calculated using CFX Maestro 2.0 (version 5.2.008.0222, Bio-Rad). Error bars represent the standard error of the mean. Statistical significance compared to *bin3^TG4.0^/+* (black asterisks) or *bin3^TG4.0^/Df; UAS-bin3/+* (sky blue asterisks) was calculated by one-way ANOVA with Benjamini-Hochberg correction for false discovery rate, using CFX Maestro 2.0 (version 5.2.008.0222, Bio-Rad). *P* values of <0.05 (*), <.01 (**), <.0001 (****), and not significant (ns) are indicated.

**Figure 6.**
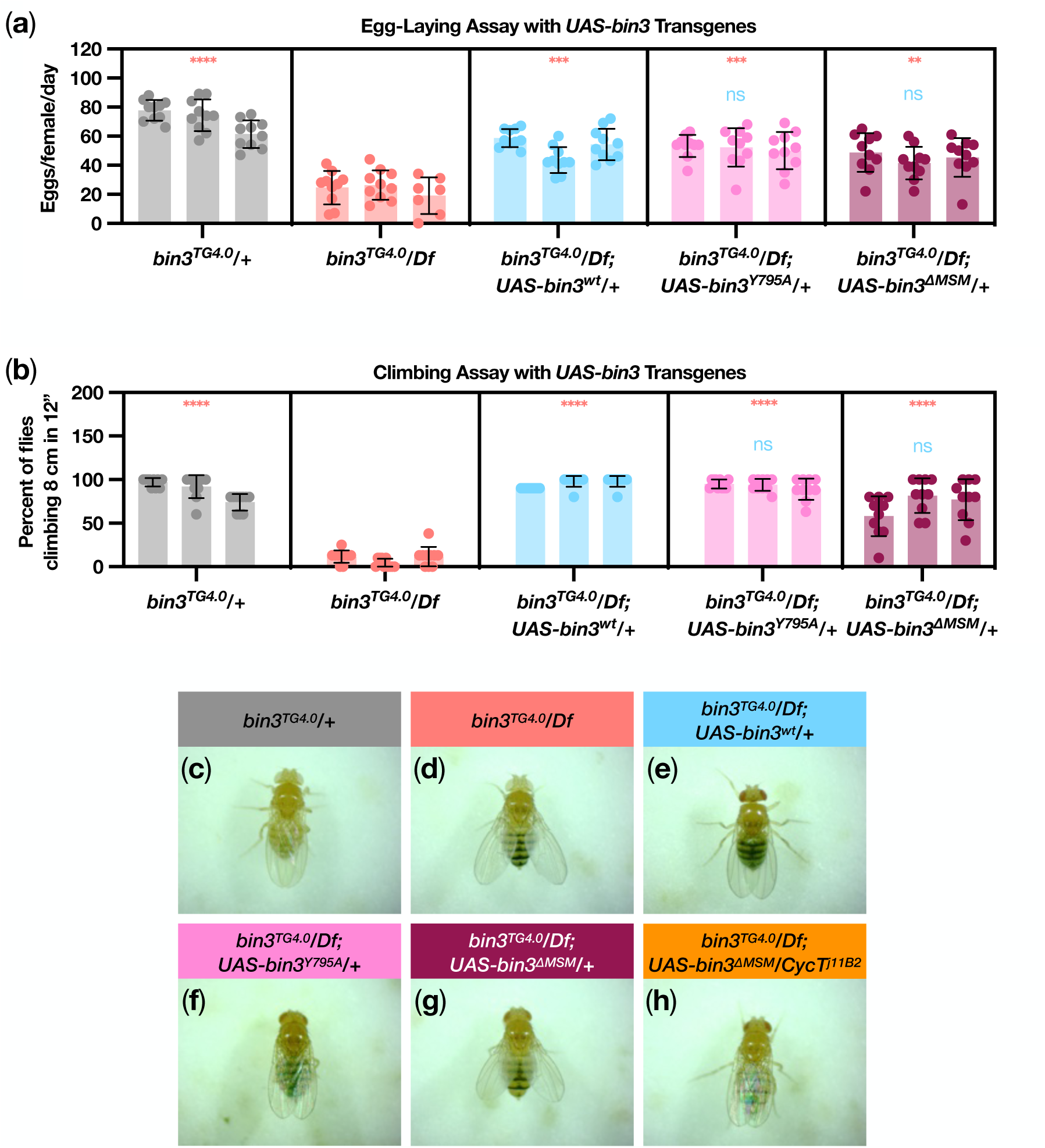
Phenotypic characterization of Bin3. ^Y795A^ **and Bin3** ^ΔMSM^ **mutant proteins *in vivo*. (a)** Scatter plot showing egg-laying rates of females of the indicated genotypes for each of three biological replicates. Error bars represent the standard deviation. Statistical significance compared to *bin3^TG4.0^/Df* (salmon asterisks) or *bin3^TG4.0^/Df; UAS-bin3/+* (sky blue asterisks) was calculated by nested one-way ANOVA with Dunnett ’s Correction for multiple comparisons, using Prism (version 9.4.1, GraphPad). *P* values of <.01 (**), <0.001 (***), <.0001 (****), and not significant (ns) are indicated. **(b)** Scatter plot showing the percent of flies of the indicated genotypes climbing 8 cm in 12” in each of 10 trials, for each of three biological replicates. Error bars represent the standard deviation. Statistical significance compared to *bin3^TG4.0^/Df* (salmon asterisks) or *bin3^TG4.0^/Df; UAS-bin3/+* (sky blue asterisks) was calculated by nested one-way ANOVA with Dunnett ’s Correction for multiple comparisons, using Prism (version 9.4.1, GraphPad). *P* values of <.0001 (****) and not significant (ns) are indicated. **(c-h)** Images of females of the indicated genotypes, exhibiting either a normal resting wing posture or a *held-out wings* phenotype.

### A metazoan-specific motif in Bin3 is required to promote specific neuromuscular functions

Much attention has been given to the methyltransferase domain of Bin3 orthologs; however, metazoans have acquired over evolutionary time additional protein sequences outside of the methyltransferase domain (Cosgrove *et al*. 2012). To determine whether there is conserved homology between Bin3 orthologs in these regions, we aligned several Bin3 orthologs that have been previously characterized, including *Drosophila* Bin3 (Zhu and Hanes 2000; Singh *et al*. 2011; Cosgrove *et al*. 2012), human MEPCE (Xue *et al*. 2010; Muniz *et al*. 2013; Shelton *et al*. 2018; Schneeberger *et al*. 2019; Yang *et al*. 2019; Lee *et al*. 2020; Zhang *et al*. 2021), zebrafish Bcdin3 (Barboric *et al*. 2009), and *S. pombe* Bmc1 (Páez-Moscoso *et al*. 2022; Porat *et al*. 2022). We discovered a 16-amino acid motif that is almost entirely conserved in metazoans (flies, zebrafish, and humans), but is absent in the fission yeast *S. pombe* (**Fig. 4c**). Therefore, we have named this motif the <u>M</u>etazoan<u>-S</u> pecific <u>M</u>otif (MSM). The N-terminal portion of this motif is predicted to be more structured (IUPred3 score 0.2733 to 0.3764), while the C-terminal portion is predicted to be less structured (IUPred3 score 0.4062 to 0.5887). Given the strong conservation of the MSM, we hypothesized that the MSM must contribute to some important function of Bin3 and its metazoan orthologs. To test this hypothesis, we used the Trojan GAL4 system to generate flies expressing either Bin3^wt^ or Bin3 deleted of the entire 16-aa MSM (Bin3^ΔMSM^) and asked whether Bin3^ΔMSM^ could (i) bind and stabilize 7SK, and (ii) rescue *bin3* mutant phenotypes.

We found that GAL4 expressed from *bin3^TG4.0^* induced both the *UAS-bin3^wt^* (**Fig. 4h**) and *UAS-bin3^ΔMSM^* (**Fig. 4j**) transgenes to express similar levels of mRNA (*P* = .3840, **Fig. 4k**) and protein (*P* = .7994, **Fig. 4l**). Given that the MSM is outside of the Bin3 methyltransferase domain, we did not expect that the Bin3 MSM would be required for 7SK stability or binding. Accordingly, there was no significant difference in 7SK levels (*P* = .3846, **Fig. 5a**) between Bin3^ΔMSM^ flies (*bin3^TG4.0^/Df; UAS-bin3^ΔMSM^/+*) and Bin3^wt^ flies (*bin3^TG4.0^/Df; UAS-bin3^wt^/+*). Also, Bin3^ΔMSM^ immunoprecipitated as much 7SK (*P* = .9938, **Fig. 5b**) as did Bin3^wt^.

Given that 7SK was bound and stabilized by Bin3 ^ΔMSM^, we expected that Bin3^ΔMSM^ would rescue the fecundity, climbing, and wing posture defects of *bin3^TG4.0^/Df* flies. Indeed, Bin3^ΔMSM^ expression rescued the fecundity (*P* = .0061, **Fig. 6a**) and climbing (*P* < .0001) defects of *bin3^TG4.0^/Df* flies, similar to Bin3^wt^ (*P* = .6681, **Fig. 6a** ; *P* = .0685, **Fig. 6b**). However, Bin3^ΔMSM^ did not rescue the *held-out wings* phenotype of *bin3^TG4.0^/Df* flies (compare **Fig. 6g** with **Fig. 6d**), suggesting that while the MSM is dispensable for Bin3 function in the contexts of egg-laying and climbing, the Bin3 MSM plays a critical role in normal resting wing posture, independent of 7SK (**Fig. 6i**). Interestingly, reducing CycT dosage in Bin3^ΔMSM^ flies (*bin3^TG4.0^/Df; UAS-bin3^ΔMSM^/CycT^j11B2^*) rescued the *held-out wings* defect (compare **Fig. 6h** with **Fig. 6g**). We think that because Bin3^ΔMSM^ binds and stabilizes 7SK (and as such, forms the 7SK snRNP to repress P-TEFb; **Fig. 5b**) it is unlikely that the MSM directly contributes to P-TEFb repression; instead, genetically reducing P-TEFb activity probably had a non-specific rescuing effect by globally tapering transcription elongation.

### Bin3 binds to the U6 spliceosomal RNA, but is not required for its stability

MEPCE in humans (Jeronimo *et al*. 2007; Muniz *et al*. 2013) and Bmc1 in*S. pombe* (Páez-Moscoso *et al*. 2022; Porat *et al*. 2022) bind to the U6 spliceosomal snRNA, but are not required for U6 stability. We asked whether Bin3 in *Drosophila* also binds to U6 and whether Bin3 is also dispensable for U6 stability.

We found that Bin3 ^wt^ (*P* = .0113), Bin3^Y795A^ (*P* = .0033), and Bin3^ΔMSM^ (*P* = .0336) all bind to U6 snRNA, compared to un-tagged control. We also found that there is no significant difference in the amount of U6 bound by Bin3^Y795A^ (*P* = .7671) or Bin3^ΔMSM^ (*P* = .8486) compared to Bin3^wt^ (**Fig. 5d**). These results show for the first time that Bin3 in *Drosophila* binds to U6, and that neither catalytic activity nor the MSM are required for binding. It is not surprising that Bin3^ΔMSM^ binds to U6, as Bmc1 lacks this region (Cosgrove *et al*. 2012), yet it binds to U6 (Páez-Moscoso *et al*. 2022; Porat *et al*. 2022). And, like MEPCE and Bmc1, we found that Bin3 is not required for U6 stability. U6 levels (**Fig. 5c**) were not different between flies expressing Bin3^wt^ and Bin3^Y795A^ (*P* = .8794) or Bin3^ΔMSM^ (*P* = .7529), and were not significantly affected in *bin3^2-7^/Df* flies (*P* = .9314, **Fig. 1d**). U6 levels were reduced by approximately half in *bin3^TG4.0^/Df* flies (*P* = .0471, **Fig. 4g**); however, these data were just below the threshold for significance (a = 0.05). Taken together, Bmc1, Bin3, and MEPCE have a conserved function in binding to the U6 snRNA, but are not required for its stability.

## DISCUSSION

In this study, we showed that Bin3, the *Drosophila* ortholog of the 7SK methylphosphate capping enzyme, MEPCE, has a conserved role in promoting normal metazoan development by regulating the P-TEFb positive elongation factor (**Figs. 7a, 8a**). We further investigated Bin3 by determining the *in vivo* requirement for two different regions of the Bin3 protein: the methyltransferase domain, and a previously undiscovered motif that is specific to metazoan orthologs of Bin3. To our surprise, we found that the catalytic activity of the methyltransferase domain appears to be dispensable for the function of Bin3 as a constituent of the repressive 7SK snRNP (**Fig. 7b**). By contrast, the metazoan-specific motif in Bin3 is required for a tissue-specific function, one that is 7SK-independent (**Figs. 7c, 8b**). We also provide the first evidence in *Drosophila* for binding of Bin3 to the U6 spliceosomal snRNA. As in other model organisms (Jeronimo *et al*. 2007; Muniz *et al*. 2013; Páez-Moscoso *et al*. 2022; Porat *et al*. 2022), we found that Bin3 was not required for U6 stability.

**Figure 7.**
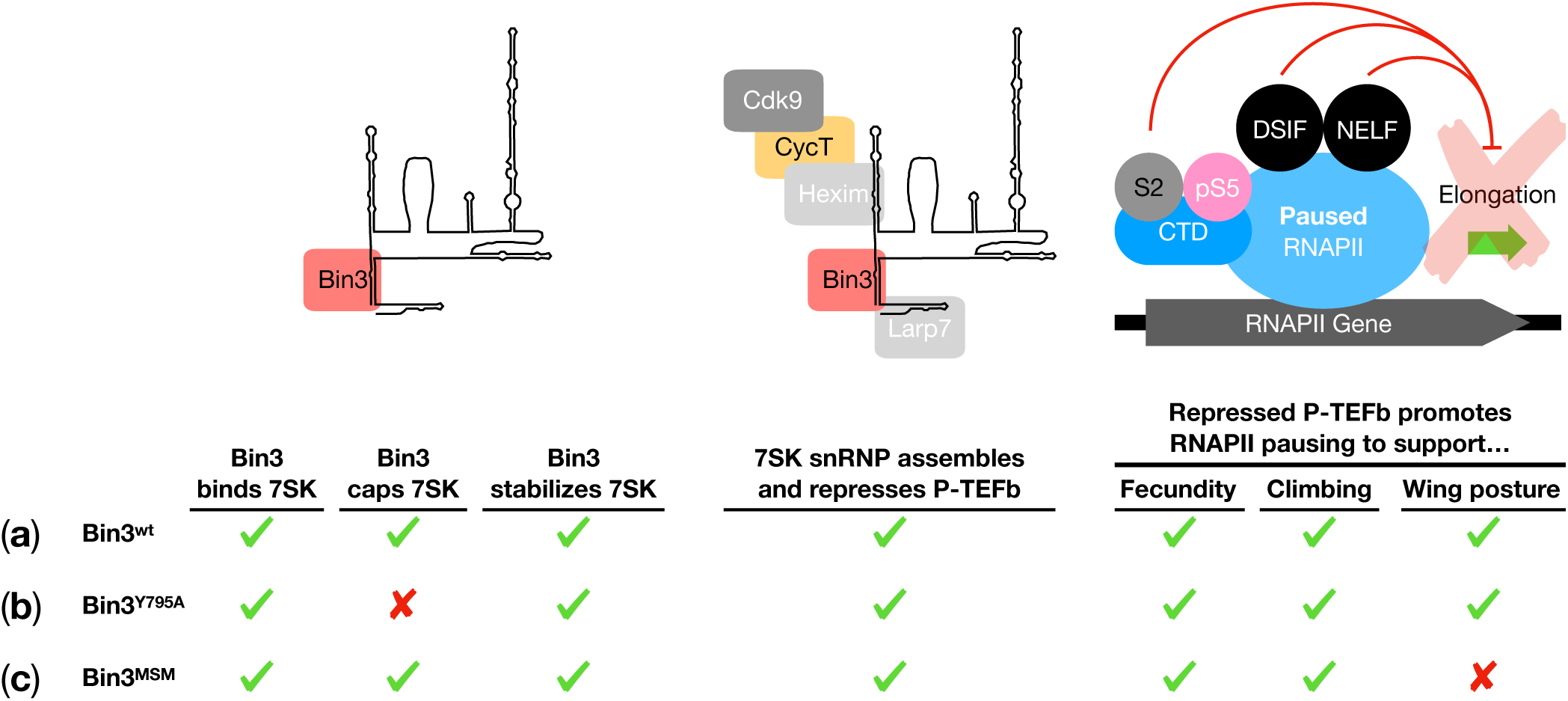
Summary of Bin3. ^wt^**, Bin3** ^Y795A^**, and Bin3** ^ΔMSM^ **protein functions. (a)** Bin3^wt^ binds, caps, and stabilizes 7SK, which allows for P-TEFb to be sequestered and repressed, which prevents RNAPII pause-release and promotes egg-laying, climbing, and normal resting wing posture. **(b)** Bin3^Y795A^ is able to bind and stabilize 7SK despite being catalytically-dead; consequently, Bin3 ^Y795A^ also promotes egg-laying, climbing, and normal resting wing posture. **(c)** Bin3^ΔMSM^ binds, caps, and stabilizes 7SK, and represses P-TEFb; this is sufficient to promote normal egg-laying and climbing, but is not sufficient to promote normal wing posture. This suggests the MSM confers a 7SK-independent, tissue-specific function to Bin3.

**Figure 8.**
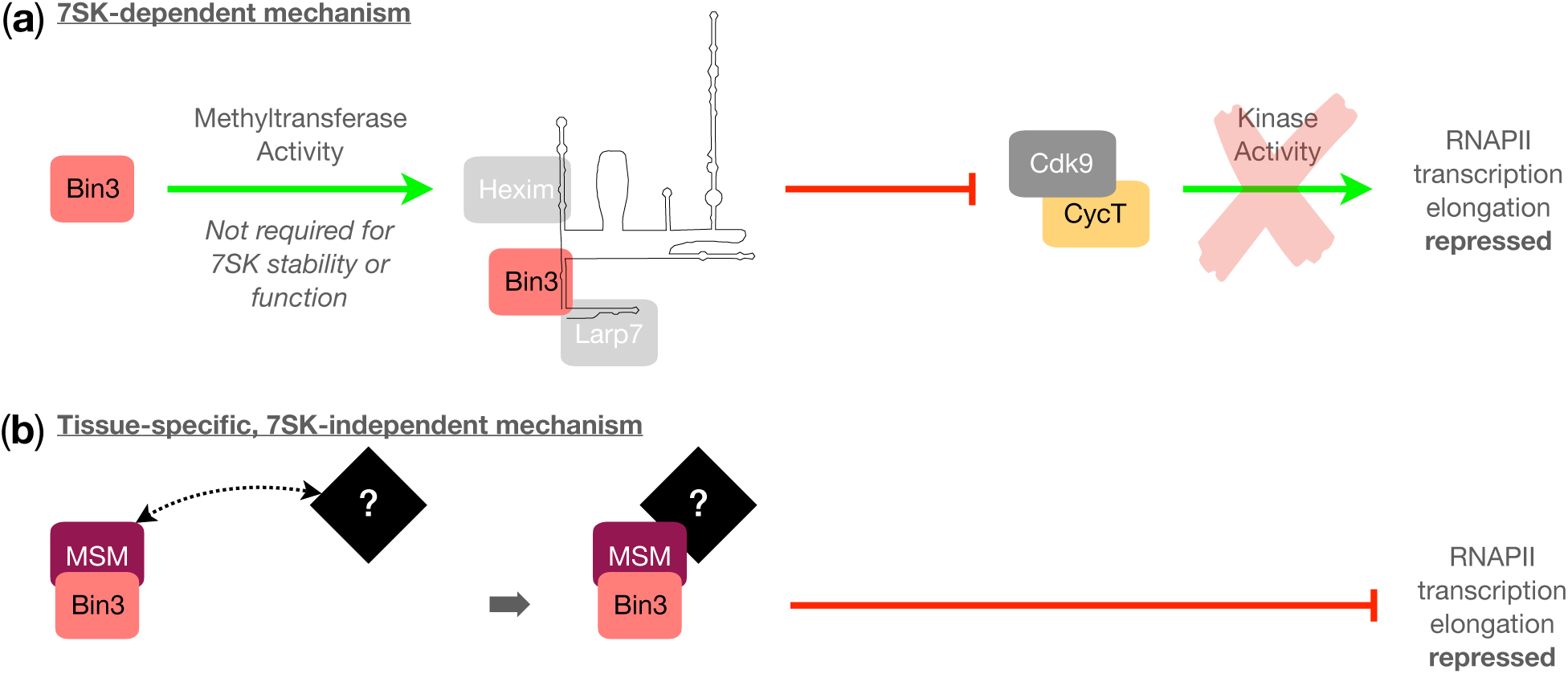
Model for the role of Bin3 in *Drosophila* development. **(a)** Bin3 binds, caps, and stabilizes 7SK; however, methyltransferase activity is not required for 7SK stability *in vivo*. Stabilized 7SK assembles a snRNP that represses P-TEFb, which prevents RNAPII transcription elongation. **(b)** The Bin3 MSM confers a 7SK-independent, tissue-specific function to Bin3. The MSM might mediate an interaction with a tissue-specific factor, or this interaction might only be important in specific tissues. Bin3 and this factor then repress transcription elongation by a mechanism that is likely downstream of P-TEFb.

### Bin3 represses P-TEFb to promote normal fly physiology

The canonical function of MEPCE is to *repress* gene expression by binding to and stabilizing 7SK snRNA, which acts as a scaffold for proteins that sequester P-TEFb (Peterlin *et al*. 2011; Cosgrove *et al*. 2012). This inhibits P-TEFb, which normally phosphorylates RNAPII to promote transcription elongation. Consequently, loss of MEPCE function destabilizes 7SK, dissociates the repressive snRNP, releasing P-TEFb (Fujinaga *et al*. 2023). Hyper-activated P-TEFb induces aberrant gene expression by ectopic phosphorylation of RNAPII and other factors (He *et al*. 2008; Schneeberger *et al*. 2019). A non-canonical function of MEPCE is to *activate* gene expression by binding to the tail of histone H4 and recruiting free P-TEFb to chromatin. This non-canonical function was found be required for the induction of tumorigenic genes in a breast cancer cell line (Shelton *et al*. 2018). Here, we show that Bin3 contributes to normal fly physiology through a mechanism most consistent with its canonical function in repression of P-TEFb. We found that the egg-laying, climbing, and wing posture defects of *bin3* mutants were rescued by genetically reducing the activity of P-TEFb with a mutation in *CycT* (which is needed for P-TEFb kinase activity). Thus, in fly development, Bin3 appears to repress P-TEFb (and gene expression) rather than to activate P-TEFb (**Figs. 7a, 8a**).

### Bin3 is required for reproductive success

Previously, we found that Bin3 plays a role in fertility by promoting embryonic viability (Singh *et al*. 2011). Here, we show that Bin3 is required for fecundity by supporting a normal egg-laying rate (**Fig. 7a**). Thus, Bin3 has a role in two different aspects of reproductive success: fertility (offspring viability), and fecundity (the rate of offspring production).

Although Bin3 and its substrate 7SK are expressed in all germline and somatic cells of the ovary, germline expression of Bin3 was dispensable for a normal egg-laying rate. A germline function for Bin3 might occur later in development, consistent with our previous finding that Bin3 appears to regulate the translation of *grk* in late-stage oocytes (Singh *et al*. 2011), which is important for embryonic patterning.

A previous report demonstrated that both wild-caught flies harboring indels in *bin3*, and transgenic flies harboring transposon insertions into *bin3*, have reduced numbers of ovarioles (Lobell *et al*. 2017). However, we found that our*bin3^2-7^/Df* mutant females did not exhibit reduced ovariole numbers compared to controls. We suggest that the difference between our findings and those of Lobell *et al*. (2017) might be attributable to differences in genetic backgrounds. Nevertheless, the wild-type number of ovarioles established in our *bin3* mutants is not sufficient to promote a normal egg-laying rate. Egg-laying is in part a neuromuscular process, as muscle contractions of the ovary and uterus promote successful oviposition (i.e., egg-laying; Deady and Sun 2015; Irizarry and Stathopoulos 2015; Liao and Nässel 2020; Garrett *et al*. 2023). It is possible, therefore, that defects in neuromuscular function that we observe in *bin3* mutants (see below) might also explain why *bin3* mutant females lay fewer eggs, *e.g.*, due to a reduction in the frequency or strength of these contractions.

### Conserved role of Bin3 and MEPCE in metazoan neurodevelopment

A patient with a neurodevelopmental disorder characterized by mobility and musculature deficiencies was shown to carry a heterozygous nonsense mutation in *MEPCE* (Schneeberger *et al*. 2019). Similarly, we found that *bin3* mutant flies exhibited difficulties in climbing and an inability to properly fold their wings, phenotypes that are characteristic of neuromuscular defects (Shukla *et al*. 2013; Kairamkonda and Nongthomba 2014; Sanhueza *et al*. 2014; Manjila and Hasan 2018). Many mutations in *Drosophila* can affect wing posture (see below), including those in the highly-conserved JAK/STAT pathway (Harrison *et al*. 1998; Callus and Mathey-Prevot 2002; Ayala-Camargo *et al*. 2013; Hatini *et al*. 2013; Johnstone *et al*. 2013). Therefore, the conserved roles of Bin3 and MEPCE in metazoan neurodevelopment may occur through the JAK/STAT pathway. While the exact mechanisms remain to be determined, the phenotypes we observe in *bin3* mutants (egg-laying, climbing, wing posture) can be attributable to neuromuscular defects, suggesting a strong conservation in the developmental function of Bin3 and MePCE in *Drosophila* and human development.

Bin3/MEPCE methylates the gamma phosphate on the 5’ end of the 7SK snRNA, while another class of RNA methyltransferases methylates the *N*^6^ position of internal adenosines (*N*^6^-methyladenosine, m^6^A) on “RRACH” motifs (Wang *et al*. 2018) in a variety of RNAs (Lence *et al*. 2018). Interestingly, both human 7SK (Warda *et al*. 2017; Leger *et al*. 2021; Perez-Pepe *et al*. 2023) and mouse 7SK (Xu *et al*. 2022) contain the m^6^A modification. In HeLa cells, methylation of 7SK by the METLL3 m^6^A “writer” protein in response to EGF signaling allows hnRNPs to associate with 7SK; this causes HEXIM to dissociate from the 7SK snRNP and release P-TEFb, which activates transcriptional elongation (Perez-Pepe *et al*. 2023). In mice, methylation of 7SK at adenine 281 by METLL3 is required for the binding of m^6^A “reader” proteins to establish proper 7SK secondary structure (Xu *et al*. 2022). *Drosophila* 7SK contains a single RRACH motif at position 397-401; however, it is not known whether the adenine within this motif is methylated, or if m ^6^A at that position would serve functions similar to those in HeLa cells or mice. Intriguingly, flies mutant for m ^6^A writers (and m^6^A readers) exhibit the same neuromuscular defects in both climbing and resting wing posture (Lence *et al*. 2016; Kan *et al*. 2017) as flies mutant for*bin3* (this study). Therefore, it is possible that *Drosophila* 7SK might also contain the m^6^A modification (at A399), and that m ^6^A is important for 7SK stability and function in neurodevelopmental processes in *Drosophila*.

Although Bin3 and the m ^6^A pathway might function synergistically to promote 7SK stability and function, they seem to act antagonistically on transcription elongation: Bin3 is required to *prevent* RNAPII pause-release, whereas the m ^6^A pathway has been shown in *Drosophila* to *promote* RNAPII pause-release (Akhtar *et al*. 2021). Clearly, there is a complex balance between 7SK stability (via Bin3 binding and possible m ^6^A methylation) as well as 7SK functions that sequester P-TEFb (Bin3-mediated function) or release P-TEFb (m ^6^A modification function) that is governed by the activities of diverse classes of RNA methyltransferases, which in *Drosophila* contribute to neuromuscular function.

Moreover, it appears that m6A modifications in 7SK might serve species-specific roles in 7SK biology. Neuromuscular disorders arise from developmental defects in motoneurons and/or the muscles that they control. We attempted to identify the cells and tissues that require Bin3 for neuromuscular function by performing an RNAi screen using over 90 GAL4 drivers active in specific neuron and muscle cell types that drive the expression of two different short hairpin RNAs (shRNAs) targeting *bin3* coding sequence (Ni *et al*. 2011). However, we did not identify any conditions under which *bin3* RNAi knockdown recapitulated the neuromuscular phenotypes of *bin3* mutant flies. The failure of our screen might be due to technical reasons such as weak GAL4 drivers and/or insufficient levels of RNAi knockdown, or they could be due to Bin3 being required in more than one tissue simultaneously for neuromuscular activity. Indeed, knockdown of *bin3* induced by the Trojan-GAL4 *bin3* allele (which also functions as a null allele) in the spatiotemporal expression pattern of *bin3* recapitulated the *held-out wings* phenotype of *bin3* mutant flies (data not shown), suggesting that Bin3 might be required in multiple cell types simultaneously, at least for normal resting wing posture.

### Bin3 catalytic activity is dispensable for 7SK stability and snRNP function *in vivo*

MEPCE modifies the 5’ end of 7SK RNA by catalyzing the addition of a monomethyl cap to the gamma phosphate (Gupta *et al*. 1990b; Shumyatsky *et al*. 1990; Jeronimo *et al*. 2007; Xue *et al*. 2010; Yang *et al*. 2019). It has long been thought that the monomethyl cap is essential for 7SK stability. For example, *in vitro* transcribed capped 7SK was more stable than uncapped 7SK when injected into *Xenopus* oocytes (Shumyatsky *et al*. 1993). However, whether 7SK requires a monomethyl cap for stability *in vivo* has never been adequately addressed. We and others have shown that Bin3 and its orthologs are essential for 7SK stability, as deletion of *bin3* (Singh *et al*. 2011), siRNA-mediated knockdown of *MEPCE* in human cells (Jeronimo *et al*. 2007; Xue *et al*. 2010), and morpholino-mediated disruption of *mepcea* splicing in zebrafish embryos (Barboric *et al*. 2009) reduces 7SK levels. Importantly, these approaches do not explicitly test the *in vivo* requirement for methyltransferase activity, which requires the use of catalytic mutants.

Catalytic mutants of MEPCE have been characterized *in vitro*. A triple mutation in the active site (V447A, L448A, and D449A) abolished methyltransferase activity, likely by preventing the interaction between MEPCE and the methyl donor *S* -adenosyl-methionine (SAM). However, this mutant is also defective for binding to 7SK and therefore it is not a catalytic mutant *per se* (Xue *et al*. 2010). A double mutant in the active site (G451A and G455A) also abolished MEPCE methyltransferase activity, again, presumably by preventing the interaction between MEPCE and SAM; however, this mutant does bind to 7SK (Shelton *et al*. 2018). Finally, a single amino acid mutation in the active site (Y421A) reduced methyltransferase activity to 0.6% that of the wt enzyme. This is likely due to loss of hydrogen bonds between Y421 and both an adjacent residue (R425) and the 7SK gamma phosphate, which is the recipient of the methyl group from SAM (Yang *et al*. 2019). Importantly, Y421A is not expected to disrupt binding to either the methyl donor co-factor or the RNA substrate, but rather to disrupt the chemistry of methyl transfer to the gamma phosphate (Yang *et al*. 2022). Therefore, given the precise and specific nature of this active site mutation, our preferred choice for *in vivo* study of the importance of methyl capping was a Y795A substitution in Bin3 (orthologous to Y421A in MEPCE).

Unexpectedly, we found that the Bin3 ^Y795A^ mutant did not exhibit any of the physiological defects exhibited by *bin3* deletion mutants, suggesting that catalytic activity is not required for Bin3 function *in vivo* (**Fig. 8a**). This result could be explained, in part, by our finding that Bin3^Y795A^ still bound and stabilized 7SK no differently than Bin3 ^wt^, despite its expected lack of methyltransferase activity (**Fig. 7b**). Thus, flies expressing the Bin3 ^Y795A^ allele retain the ability to assemble the 7SK snRNP to repress P-TEFb and promote normal development. Our findings suggest that *in vivo* capping of endogenous 7SK is not required for its stability *per se*, so long as Bin3 is able to bind to 7SK.

That the Bin3 ^Y795A^ catalytic mutant protein was still able to bind to 7SK is not entirely surprising. Y421 in MEPCE (analogous to Y795 in Bin3) is located within a disordered region that becomes ordered upon binding to 7SK, forming an alpha helix (Yang *et al*. 2022); this disorder-to-order transition is required for MEPCE methyltransferase activity. When LARP7 binds to 7SK and then associates with MEPCE, the alpha helix containing Y421 becomes disordered again (Yang *et al*. 2022); this order-to-disorder transition induced by LARP7 binding inhibits MEPCE methyltransferase activity (Xue *et al*. 2010; Brogie and Price 2017; Yang *et al*. 2022). Therefore, MEPCE bound by LARP7 effectively phenocopies MEPCE with a Y421A mutation, which is catalytically-dead (Yang *et al*. 2019). Thus, it appears that Y421 in MEPCE and Y795 in Bin3 are not essential for 7SK binding, but instead are essential only for methyl capping.

Our finding that the Bin3 ^Y795A^ catalytic mutant protein not only bound 7SK, but stabilized 7SK despite the expected lack of catalytic activity, suggests that RNA binding contributes more to 7SK stability than does methyl capping, challenging the assumption that the gamma monomethyl cap is essential for 7SK stability. This is underscored by the fact that U6 snRNA, which binds to and is presumed to be capped by Bin3 and its orthologs, is stable in the absence of these proteins (Jeronimo *et al*. 2007; Muniz *et al*. 2013; Páez-Moscoso *et al*. 2022; Porat *et al*. 2022).

Bin3 and its orthologs are peculiar enzymes, in that they remain associated with the products of the reaction they catalyze (Xue *et al*. 2010; Yang *et al*. 2019, 2022). MEPCE has a higher affinity both for capped 7SK and SAH (the products) than uncapped 7SK and SAM (the reactants) (Yang *et al*. 2019). Therefore, the gamma monomethyl cap might be required not for 7SK stability *per se*, but to enhance the binding of MEPCE to the 5’ end of 7SK to keep the enzyme bound after catalyzing the methyltransferase reaction. Constitutive binding of MEPCE to 7SK would serve the dual purposes of physically protecting 7SK from inorganic phosphatases (for which the gamma monomethyl cap is a substrate; Shumyatsky *et al*. 1990), and aiding in the recruitment of LARP7, which binds weakly to 7SK in the absence of MEPCE (Xue *et al*. 2010), but enhances the affinity of MEPCE for 7SK after binding (Muniz *et al*. 2013; Yang *et al*. 2022). However, our finding that the Bin3^Y795A^ catalytic mutant bound as much 7SK as Bin3 ^wt^ despite the lack of a gamma monomethyl cap is not consistent with such a model (at least not *in vivo*). Instead, our results support a model in which 7SK RNA secondary structure—and not the gamma monomethyl cap—is required for Bin3 and its orthologs to bind to 7SK. Indeed, MEPCE bound equally well to transfected 7SK harboring either the normal gamma monomethyl cap or a tri-methyl guanosine (TMG) cap (Muniz *et al*. 2013), and Bmc1 (the *S. pombe* ortholog of Bin3) binds to both gamma monomethyl-capped U6 RNA and TMG-capped telomerase RNA (Páez-Moscoso *et al*. 2022; Porat *et al*. 2022). These findings suggest that a gamma monomethyl cap structure is not specifically required for constitutive or high-affinity binding by MEPCE and its orthologs. Alternatively, 7SK secondary structure features likely mediate initial association with MEPCE (i.e., prior to the association of LARP7, which enhances the affinity of MEPCE for 7SK). When MEPCE initially binds to 7SK in the “nascent” conformation (adopted during 7SK synthesis), there is extensive hydrogen bonding between the methyltransferase domain and the 5’ hairpin of 7SK (Yang *et al*. 2019). Binding of MEPCE to 7SK requires the first two (Singh *et al*. 1990) to three (Muniz *et al*. 2013) base pairs of the 5’ hairpin, and a portion of the single-stranded DNA 3’ to the hairpin (Shumyatsky *et al*. 1994; Muniz *et al*. 2013; Yang *et al*. 2019). Subsequently, LARP7 binds to the complex of MEPCE and 7SK, and the conformation of 7SK changes from a “nascent” conformation to a “closed” conformation (adopted after formation of the core snRNP of MEPCE, 7SK, and LARP7). In this conformation, MEPCE forms additional hydrogen bonds to 7SK, and thereby has higher affinity for the “closed” conformation of 7SK than for the “nascent” conformation of 7SK (Yang *et al*. 2022). Therefore, both 7SK secondary structure and association with LARP7 are likely the major contributors to the constitutive binding of MEPCE to 7SK.

One possible circumstance under which the gamma monomethyl cap on 7SK might be essential for stability is when the 7SK snRNP is actively repressing P-TEFb. For HEXIM to bind to 7SK and sequester P-TEFb, 7SK must undergo a conformational change from the “closed” conformation (i.e., the “core” snRNP conformation) to the “linear” conformation (i.e., the “repressive” snRNP conformation). In this conformation, the 5’ end of 7SK is located outside of the MEPCE active site (Yang *et al*. 2022), and therefore, in the context of the repressive 7SK snRNP, the 5’ end of 7SK might be solvent-exposed, which would necessitate a gamma monomethyl cap to protect 7SK from exoribonucleolytic degradation. However, these findings are based on crystal structures using a recombinant MEPCE comprising only the methyltransferase domain. Our finding that 7SK is not only bound but is also stabilized by Bin3^Y795A^ catalytic mutant protein (just like Bin3^wt^ protein) leaves open the possibility that there are regions outside of the methyltransferase domain in full-length Bin3/MEPCE that protects the 5’ end of 7SK, even when located outside of the active site. Moreover, the finding that the 5’ end of 7SK might not be bound by MEPCE when in the “linear” conformation further supports a model in which the gamma monomethyl cap is not required for constitutive binding of MEPCE to 7SK.

If the monomethyl cap is not required for either 7SK stability or for Bin3 binding, then what is the purpose of the methyltransferase domain in Bin3 and its orthologs? One possibility is that the methyltransferase domain is a vestige of an ancestral ortholog of Bin3 that did not remain constitutively bound to its RNA substrates, and therefore, the monomethyl cap would have been essential to protect these RNAs from 5’-3’ exoribonucleolytic degradation. The acquisition of additional protein binding constituents during evolution of a primordial 7SK snRNP complex might have necessitated that Bin3 remain associated with 7SK (after catalyzing the methyltransferase reaction) to facilitate the stable interaction of these additional components (like Larp7). This process would have rendered the monomethyl capping reaction dispensable for 7SK stability, as we observe. A second possibility, supported by the strong conservation of the methyltransferase domain over evolutionary time, argues that catalytic activity has an important function that has yet to be identified, perhaps on substrates other than 7SK.

### A metazoan-specific motif in Bin3 facilitates a 7SK-independent, tissue-specific function in neurodevelopment

*S. pombe* Bmc1, the most ancient ortholog of Bin3 studied to date, consists only of the conserved methyltransferase domain (Cosgrove *et al*. 2012; Páez-Moscoso *et al*. 2022; Porat *et al*. 2022). Over the course of evolution, Bin3 orthologs gained additional protein sequence flanking the methyltransferase domain (Cosgrove *et al*. 2012). We found a 16-amino acid motif N-terminal to the methyltransferase domain that is highly conserved in metazoans, and is mostly unstructured. Given its location outside of the methyltransferase domain, we were not surprised to find that this <u>m</u> etazoan-<u>s</u>pecific <u>m</u>otif (MSM) was not required for Bin3 to bind or stabilize 7SK *in vivo*, and consequently, was dispensable for fecundity and climbing ability. What we did not expect, however, was that flies expressing Bin3 deleted of the MSM (Bin3^ΔMSM^) were unable to properly fold their wings, phenocopying *bin3* null mutants (**Fig. 7c**). These findings suggest that the Bin3 MSM plays a tissue-specific role in gene regulation independent of 7SK (**Fig. 8b**). There is likely one or more MSM-interacting proteins that help Bin3 regulate gene expression. These proteins might be specifically expressed in the tissues involved in wing posture (*e.g*., the direct and indirect flight muscles), or they might be ubiquitously expressed, but the interaction with the Bin3 MSM is essential for Bin3 function only in tissues contributing to normal resting wing posture. Moreover, this protein might be incorporated into the core snRNP of 7SK, Bin3, and Larp7 to generate a novel 7SK snRNP, or the association with Bin3 MSM might be independent of the 7SK snRNP altogether.

A 7SK-independent function of Bin3 is not without precedent. It was previously shown that fibroblasts from a patient with a nonsense mutation in *MEPCE* (*MEPCE* haploinsufficient cells) and cells deleted of *LARP7* (*LARP7* KO cells) both had reduced levels of 7SK, as expected, but that genes up-regulated in *MEPCE* haploinsufficient cells were not affected in *LARP7* KO cells. Moreover, exogenous *MEPCE* rescued the up-regulation phenotype of *MEPCE* haploinsufficient cells, despite only leading to a small increase in 7SK levels. Based on these findings, Schneeberger *et al*. (2019) hypothesized that MEPCE might have a 7SK-independent role in regulating transcription, potentially by interacting with as-yet unknown proteins. Our finding that Bin3 ^ΔMSM^ binds and stabilizes 7SK but is not sufficient to rescue all *bin3* mutant phenotypes supports this hypothesis and suggests that the as-yet unknown proteins that interact with Bin3/MEPCE to regulate transcription does so through the MSM, independent of the 7SK snRNP.

## Conclusions

In summary, we have carried out the first detailed *in vivo* study of the highly-conserved Bin3/MEPCE RNA methyltransferase in a metazoan host. The surprising results demonstrate the value of rigorously testing models for enzyme function in whole organisms. Our study establishes *Drosophila* as a genetically tractable developmental model for further dissection of Bin3/MEPCE and 7SK non-coding RNA pathways, and how their disruption may be associated with human disease.

## DATA AVAILABILITY

Plasmids and *Drosophila* stocks are available upon request. All reagent inquiries should be directed to RJP. All data required to confirm the conclusions made in this article are presented in the text, figures, and tables.

## ACKNOWLEDGEMENTS

We are grateful to the Bloomington *Drosophila* Stock Center (NIH Grant P40OD018537) for fly stocks, the *Drosophila* Genomics Resource Center (NIH Grant 2P40OD010949) for plasmid pUASP-attB; Jian-Quan Ni for plasmid pNP; BestGene, Inc. for generating transgenic flies; and David Price for anti-CycT antiserum. We are also grateful to Scott Neal, Mark Bayfield, and Jennifer Porat for helpful discussions.

## SUPPLEMENTAL TABLES

**Table S1. Public fly stocks used in this study.** This table lists the publicly available fly stocks used in this study, including their genotypes as they appear in the paper, their detailed genotypes, the stock center from which they were procured, and the stock number.

**Table S2. Fly stocks produced for this study.** This table lists the fly stocks produced for this study, including their genotypes as they appear in the paper, their detailed genotypes, and a detailed description of how each fly stock was produced.

**Table S3. Flies used for experiments.** This table lists the flies used in each figure, including their genotypes as they appear in the paper, their detailed genotypes, the genotypes of the female and male flies crossed to produce flies used for experiments, and the figure(s) in which each fly is used.

**Table S4. DNA sequences.** This table lists the names, sequences, and purposes of DNA primers and synthetic DNA constructs used for PCR, RT-PCR, RT-qPCR, and NEB Hi-Fi DNA Assembly.

